# Presynaptic and Postsynaptic Determinants of the Functional Connectivity Between the Claustrum and Anterior Cingulate Cortex

**DOI:** 10.1101/2023.03.23.533767

**Authors:** Roberto de la Torre-Martínez, Zach Chia, Anna Tokarska, Johanna Frost-Nylén, George J Augustine, Gilad Silberberg

## Abstract

The claustrum (CLA) is a brain nucleus located between the insula and lateral striatum, implicated in a wide range of behaviors. Underpinning the different behavioral phenotypes is the connectivity between the claustrum and various cortical regions, including the anterior cingulate cortex (ACC). CLA projection neurons are glutamatergic neurons, however, the impact of CLA on its cortical targets has been shown in some studies to be inhibitory. Such inhibition is likely to arise from claustral activation of cortical interneurons, however, the intricate synaptic connectivity between different CLA and cortical cell types is not known. Here, we combine *in vivo* and *ex vivo* electrophysiology and optogenetics to reveal the functional organization of the CLA-ACC pathway according to the identity of its pre- and postsynaptic populations. Optogenetic stimulation of CLA neurons in awake mice resulted in multiphasic excitatory and inhibitory responses in ACC cells, which depended on the layer, cell type, and stimulated CLA population. Using *ex vivo* paired recordings in ACC, monosynaptic responses were recorded in pyramidal cells and different types of interneurons following photostimulation of CLA-ACC synaptic terminals. CLA axons formed monosynaptic connections in all ACC cortical layers, but the probability and strength of synaptic responses depended on the type of CLA projection, target layer in ACC, and the type of postsynaptic neuron. This intricate organization of the CLA-ACC pathway may explain the complex impact of CLA on ACC and other cortical regions, thus resolving some of the discrepancies in the field and shedding light on the functional role CLA plays in cortical function.

## Introduction

The claustrum (CLA) is a highly connected brain region between the insula and striatum. Its strong reciprocal connectivity with cortical^1–21^ and subcortical^9, 22–27^ brain regions has been the impetus behind many hypotheses on its function^10, 28–36^. Functions implicated have included: awareness^37^, consciousness^38^, salience reporting^10, 11, 39–41^, memory consolidation^42^, pain mediation^35, 36^, attention^43–45^, regulation of cortical sleep waves^46–50^ and impulse control^51^. Central to the different functions assumed for CLA is the robust connectivity with various cortical regions^1, 11, 21^, in particular with prefrontal regions such as the anterior cingulate cortex (ACC)^11, 18, 20, 21^. Several studies have shown that despite being a glutamatergic excitatory projection, the impact of CLA on targeted cortical regions is largely inhibitory, mediated by synchronized activation of cortical GABAergic interneurons ^13, 42, 43, 50, 52–56^. Recent work, however, showed that projections from VGLUT2-expressing CLA neurons increased cortical activity^45^, suggesting that the impact of CLA on its cortical targets may depend on the type of projection neurons. Indeed, CLA neurons are heterogenous, containing subtypes expressing different electrophysiological properties^10, 19, 21, 24^, even among CLA neurons projecting to the same region, including ACC^10, 11^. Another factor that may contribute the complex impact of CLA on its cortical targets is the specific connectivity with different types of cortical neurons. The cortex is made up of a majority of excitatory neurons and a small but diverse population of GABAergic interneurons^57, 58^ which are instrumental in shaping cortical activity^59–64^.

Here, we built on our previous studies of the CLA-to-ACC circuit^10, 11^ and investigated the ACC targets of CLA projections. We combined *in vivo* multiunit recordings in awake mice, *ex vivo* whole-cell recordings, optogenetics, and pharmacology to investigate he functional organization of the CLA-ACC pathway. We show that CLA stimulation in awake mice produces diverse responses in the ACC with both excitation and inhibition, depending on the identity of the presynaptic CLA projection. Further delineation using *ex vivo* whole-cell recordings revealed that the CLA provided monosynaptic, excitatory input to all layers of the ACC. All major GABAergic interneuron populations and pyramidal neurons received CLA input with layer and cell type-dependent specificities. Our results enabled us to produce a schematic of CLA targets at the ACC. This first architectural plan of functional claustro-cortical targets lays the framework to understanding the detailed mechanism of CLA action on the cortex.

## Results

### CLA modulates cortical activity in the ACC in vivo

To understand the overall effect of CLA projections on the ACC, we first investigated local field potentials (LFP) and how cortical activity in the ACC layers 2/3 and 5/6 were modulated by CLA stimulation (Fig. 1). To that end, awake, head-fixed restrained mice were implanted with an optic fiber above the CLA for photostimulation and silicon probes were placed in the ACC, ipsilateral to the optic fiber, to record the resulting postsynaptic responses (Fig. 1a, b, c, d). Both, fiber and implants were tagged with colored dyes to enable their localization (Fig. 1e, f). Prior to implantation, anterograde ChR2 virus driven by either a broad-acting CaMKIIa promoter or a specific Cre-dependent promoter was injected unilaterally into the CLA of transgenic-reporter mouse lines and the VGLUT2-Cre mouse line respectively (Fig. 1c, e, f). After at least 21 days of incubation, the mice were prepared for *in vivo* experiments.

**Fig.1.**
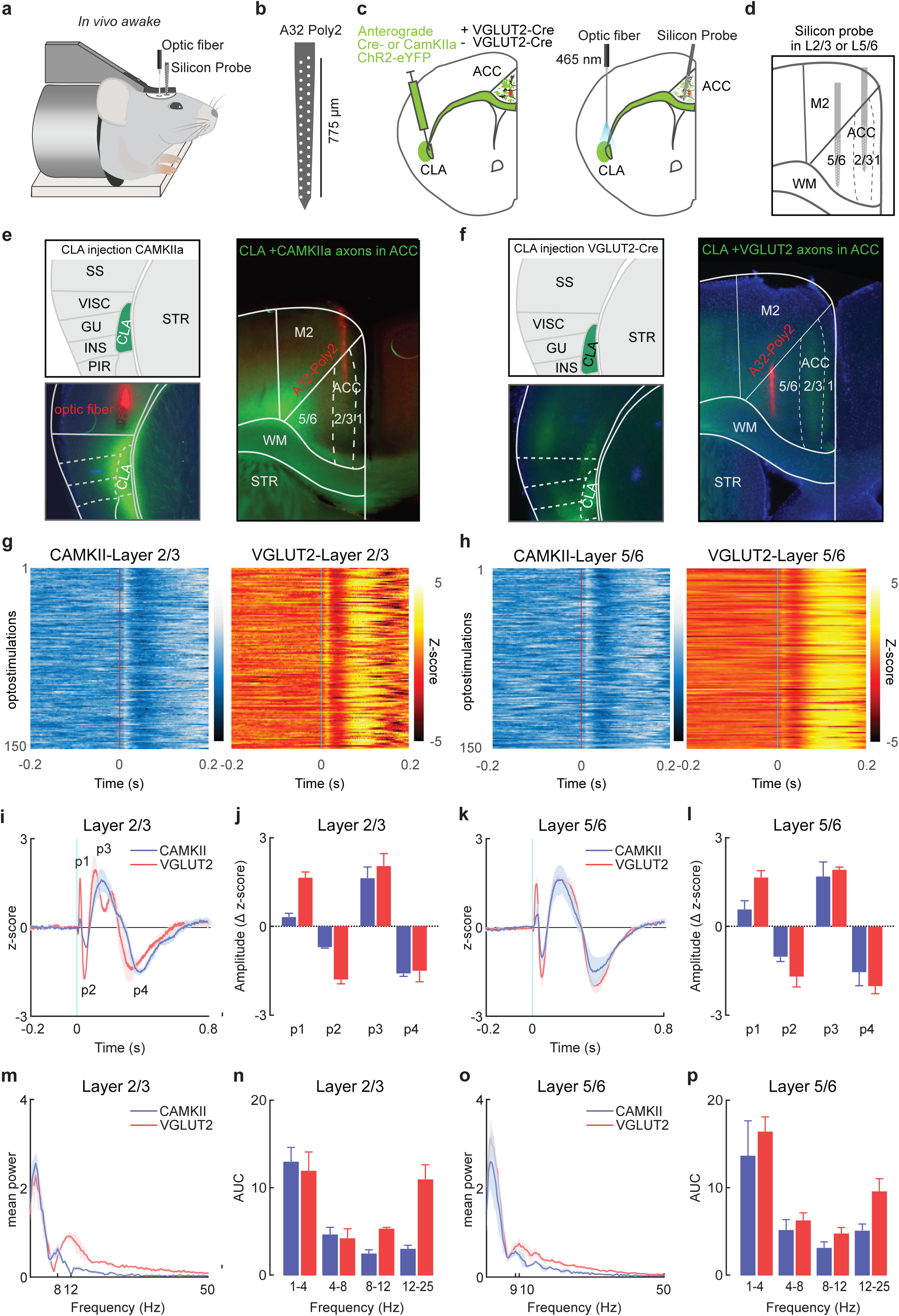
Activation of CLA-ACC neurons affects cortical activity in the ACC *in vivo*. **a.** *In vivo* recordings were performed in the mouse ACC with high-density silicon probes. **b**. Schematic of the 32 channels silicon probe (NeuroNexus A32-Poly 2). **c**. Illustration depicting experimental paradigm. Briefly, ChR2 driven by CaMKIIa promoter (-Vglu2-Cre) was injected using an anterograde viral vector into the claustrum into different transgenic mouse lines with fluorescent reporters to identify interneuron populations. For VGLUT2-Cre mice, ChR2 was injected using an anterograde Cre-dependent virus. On the day of the recording, an optic fiber was implanted on top of the CLA. **d**. Electrophysiological recordings were performed in layer 2/3 or layer 5/6. **e**. Confocal image of a 250 μm experimental slice in a PV-tdTomato mouse, with a close-up in the CLA (left panel) and ACC region (right panel) showing YFP expression in the CLA and fibers in the ipsilateral ACC but not contralateral ACC. In red, the optic fiber (left panel) and the silicon probe (right panel) are indicated. **f**. Confocal image of a 250 μm experimental slice in a VGLUT2 mouse, with a close-up in the CLA (left panel) and ACC region (right panel) showing YFP expression in the CLA and fibers in the ipsilateral ACC but not contralateral ACC. **g**. Heat map showing 150 repetitions of LFP responses in the ACC layer 2/3 to 5 ms photostimulation of CLA neurons driven by a CaMKIIa promoter (left panel) or VGLUT2 subpopulation (left panel). **h**. Same as in **g** but for layer 5/6. **i**. Grand average of the LFP responses in the ACC layer 2/3 to 5 ms photostimulation of CLA neurons driven by a CaMKIIa promoter (blue trace) or VGLUT2 subpopulation (red trace). **j**. The different excitation and inhibition are marked as p1, p2, p3, and p4. **k**. Same as in **i** but for layer 5/6. **l**. Same as in **j** but for layer 5/6. **m**. Power spectrum analysis and **n**. the area under the curve (AUC) of the oscillations evoked in the ACC after 5 ms photostimulation of CLA neurons driven by a CaMKIIa promoter (blue trace) or VGLUT2 subpopulation (red trace). **o** and **p**. Same as in **m** and **n** but for layer 5/6.

A 5 ms pulse was used to photostimulate CLA neurons, which evoked consistent triphasic LFP responses composed of an early excitation, followed by inhibition and a late excitation in both the CaMKIIa group and VGLUT2 population (Fig 1g, h). Analysis of the responses in layer 2/3 revealed that the early excitatory and inhibitory responses were larger in amplitude in the VGLUT2 population than in the CaMKIIa group (Fig 1j, l, CaMKIIa_p1_ in 2/3 = 0.3, VGLUT2_p1_ in 2/3 = 1.6, *P* < 0.05, t-test; CaMKIIa_p2_ in 2/3 = −0.7, VGLUT2_p2_ in 2/3 = −1.8, *P* < 0.05, t-test). These results were also consistent in layer 5/6, where an early excitation and inhibition component with larger amplitude in the VGLUT2 population than in the CaMKIIa group was observed (Fig 1j, l, CaMKIIa_p1_ in 5/6 = 0.5, VGLUT2_p1_ in 2/3 = 1.6, *P* < 0.05; CaMKIIa_p2_ in 2/3 = −1.0, VGLUT2_p2_ in 2/3 = −1.7, *P* < 0.05). In addition, stimulation of the VGLUT2 population (Supplementary Fig. 1) produced biphasic responses in the late excitation in layer 2/3 with an increase in alpha band power (8-12 Hz) that was not observed in either in layer 5/6 nor after broad activation using the CaMKIIa promoter (Fig 1i,k,m,n,o,p). To control for possible electrical artifacts produced by photostimulation, control experiments were performed in mice where the virus was not injected or where the virus was injected out of the CLA. These experiments revealed the absence of LFP responses in the ACC post-CLA stimulation, thus discarding a possible contribution of electrical artifacts to the LFP signal (Supplementary Fig. 2).

Taken together, our results demonstrate that projections from the CLA to the ACC are involved in modulating the excitatory and inhibitory balance in the ACC and the modulatory effect appears to be stronger in the VGLUT2 population compared to the broad activation with the CaMKIIa promoter.

### CLA activation modulates ACC preferentially affecting inhibitory interneurons in vivo

To elucidate the nature of ACC responses at a single unit level, we performed spike sorting on recordings obtained from the ACC using Kilosort2 which recovered hundreds of well-isolated units (CaMKIIa: layer 2/3 n=311 units, layer 5/6 n=77 units; VGLUT2: layer 2/3 n=142 units, layer 5/6 n=68 units). Following manual curation using Phy2 and upon examination of their waveforms, two main groups of neurons were defined: fast spiking neurons (FS), characterized by narrow spike widths (<0.5 ms), and regular spiking neurons (RS), characterized by wider spike widths (>0.7 ms) (Fig. 2b and c). Previous research has indicated that FS neurons are primarily composed of PV interneurons, while RS neurons are likely to be a heterogeneous group that includes mostly pyramidal neurons, as well as other types of interneurons^65^.

**Fig. 2.**
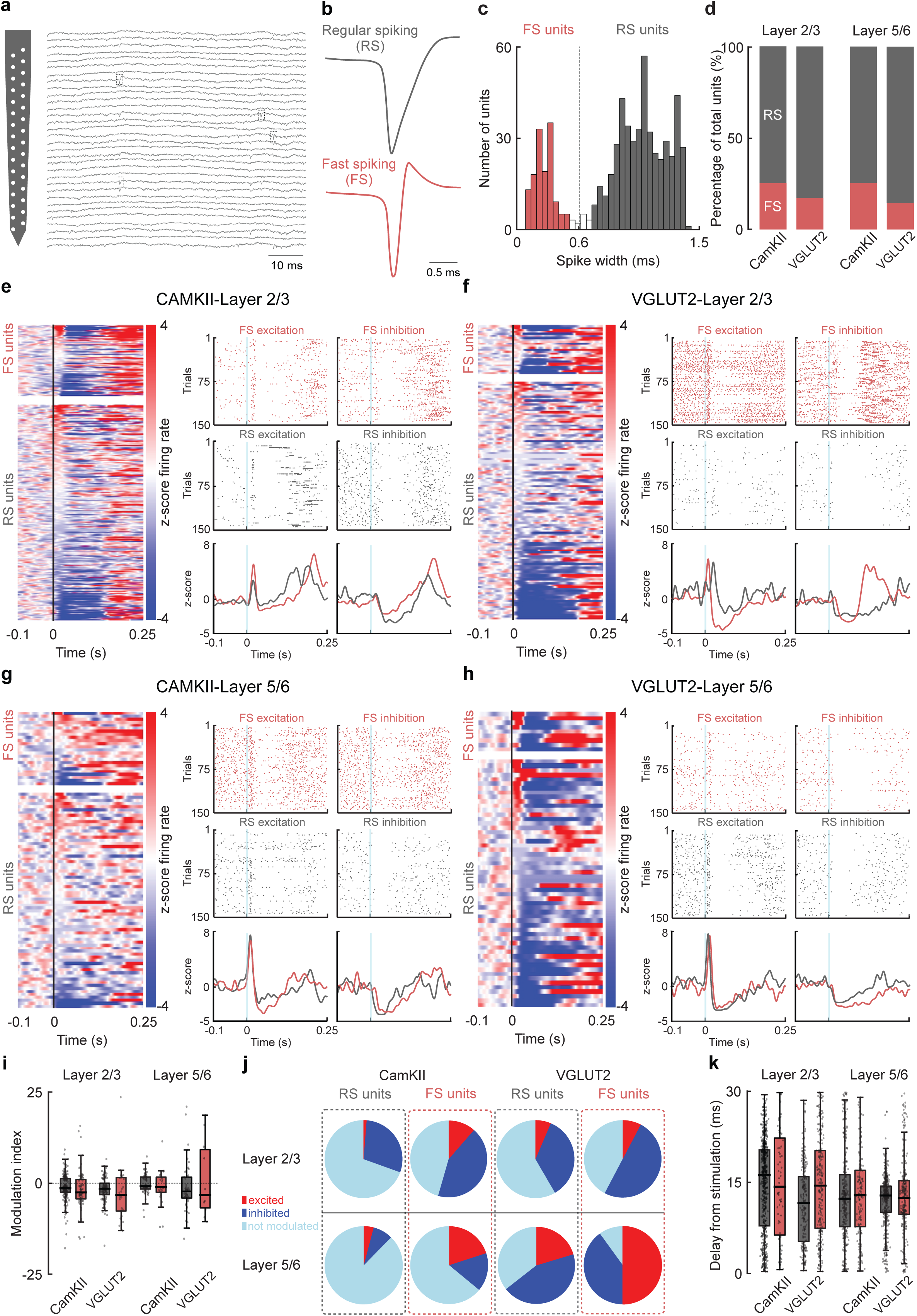
ACC cells can be excited or inhibited by broad CLA neurons activation or selective VGLUT2 CLA neurons excitation. **a.** Schematic of the 32 channels silicon probe (NeuroNexus A32-Poly 2). Representative *in vivo* recording performed in the mouse ACC with high-density 32 channels silicon probes **b**. Examples of RS (gray trace) and FS (red trace) well-isolated units in the ACC. **c**. Histogram of the distribution of the spike width of all the well-identify units in the ACC. **d**. Percentage of RS (gray bar) and FS (red bar) for each group (CaMKIIa and VGLUT2) in the different layers. **e**. Heat map showing the responses of all the units recorded in the ACC layer 2/3 to 5 ms photostimulation of CLA neurons driven by a CaMKIIa promoter VGLUT2 subpopulation (left panel). Raster plots of examples units showing excitation (left panels) or inhibition (right panels) for FS and RS. **f**. Same as in e but for VGLUT2. **g**. Same as **e** but for layer 5/6. **h**. Same as **f** but for layer 5/6. **i**. Modulation index showing the maximum or minimum z-score value in the 50 ms window after photostimulation of the CLA neurons. **j**. Proportion of RS and FS that are excited (red), inhibited (dark blue) or that were not modulated (light blue) in the ACC after photostimulation of the CLA neurons. **k**. Time delay from stimulation to maximum z-score value z-score value in the 50 ms window after photostimulation of the CLA neurons.

To evaluate whether CaMKIIa and VGLUT2 populations recruit FS and RS neurons to similar extents, we analyzed the proportion of units in each group in the CaMKIIa and VGLUT2 populations. Specifically, in layer 2/3, 25.4% of units obtained from the CaMKIIa group were classified as FS, while the proportion was slightly smaller (17.3%) in the VGLUT2 group. This trend was maintained in layer 5/6 where more FS were recorded in the CaMKIIa group than in the VGLUT2 group, although these differences were not statistically significant (Fig. 2d).

To investigate the effect of CLA neurons at the ACC, we photostimulated CLA neurons with a 5 ms pulse and observed a variety of responses at the single unit level in both the CaMKIIa and VGLUT2 populations (Fig 2e, f, g, h; trial-averaged responses of all units and example units). In layer 2/3 (Fig. 2e, f, j), activation of the CLA-CaMKIIa neurons produced excitation mainly in FS (7/85; 8.40%), with only two RS unit excited (2/226; 0.88%), but most neurons were either inhibited (FS=31/85; 35.29%; RS=38/226; 16.81%) or not affected (FS=48/85; 56.47%; RS=186/226; 82.30%). Photostimulation of the CLA-VGLUT2 population resulted in a similar proportion of excited FS units (2/24; 8.33%) and a slightly higher proportion of excited RS units (5/118; 4.23%) in layer 2/3, along with a high number of inhibited (FS=10/24; 41.67%; RS=36/118; 30.51%) or unaffected neurons (FS=12/24; 50.00%; RS=77/118; 65.05%). Overall, our results suggest that both broad and selective activation of CLA neurons leads to a negative modulation of the ACC neuronal population.

In layer 5/6 (Fig. 2g, h, j), photostimulation of CaMKIIa-expressing CLA neurons evoked a higher proportion of excitatory responses in both FS and RS units compared to layer 2/3 (FS=2/19; 10.53%; RS=3/58; 5.17%), and a lower number of inhibited units of both FS and RS populations (FS=4/19; 21.05%; RS=5/58; 8.26%). Photostimulation of VGLUT2-expressing CLA population in layer 5/6 produced stronger modulatory effects compared to layer 2/3 where just under half of the FS recorded were excited and the other half inhibited (FS_exc_ =5/11; 45.45%; FS_inh_=4/11; 36.36%). A higher fraction of RS units was excited compared to layer 2/3 and a similar proportion of units inhibited (RS_exc_ =5/57; 8.77%; RS_inh_=33/57; 57.89%). In all cases, the units were excited within a range of 12 to 15 ms making it difficult to conclude whether some of this excitation was evoked from the disinhibition of microcircuits (Fig. 2k). Taken together, our data suggests that activation of CLA neurons results in complex excitation-inhibition modulation profile of the ACC neuronal population, with a predominance of inhibition in FS and RS cells, and excitation mainly seen in FS cells.

### Claustrum provides monosynaptic input to all ACC layers

To pinpoint the cellular connectivity of CLA projections to the ACC, *ex vivo* paired whole-cell recordings were performed. Similar to *in vivo* experiments, anterograde ChR2 virus driven by a CaMKIIa promoter was injected into the CLA in different transgenic mouse lines and cortical slices were obtained from these mice after at least 21 days after viral injection. Paired recordings of a molecularly defined interneurons and neighboring pyramidal cells within the same layer were made (Fig. 3a). Although pressure injections into the CLA had minor spillage into the nearby insula (INS, Fig. 3b), this did not affect interpretation as it has been shown with retrograde beads and virus that projections to the ACC arise mainly from the CLA rather than the INS^10, 11^.

**Fig. 3.**
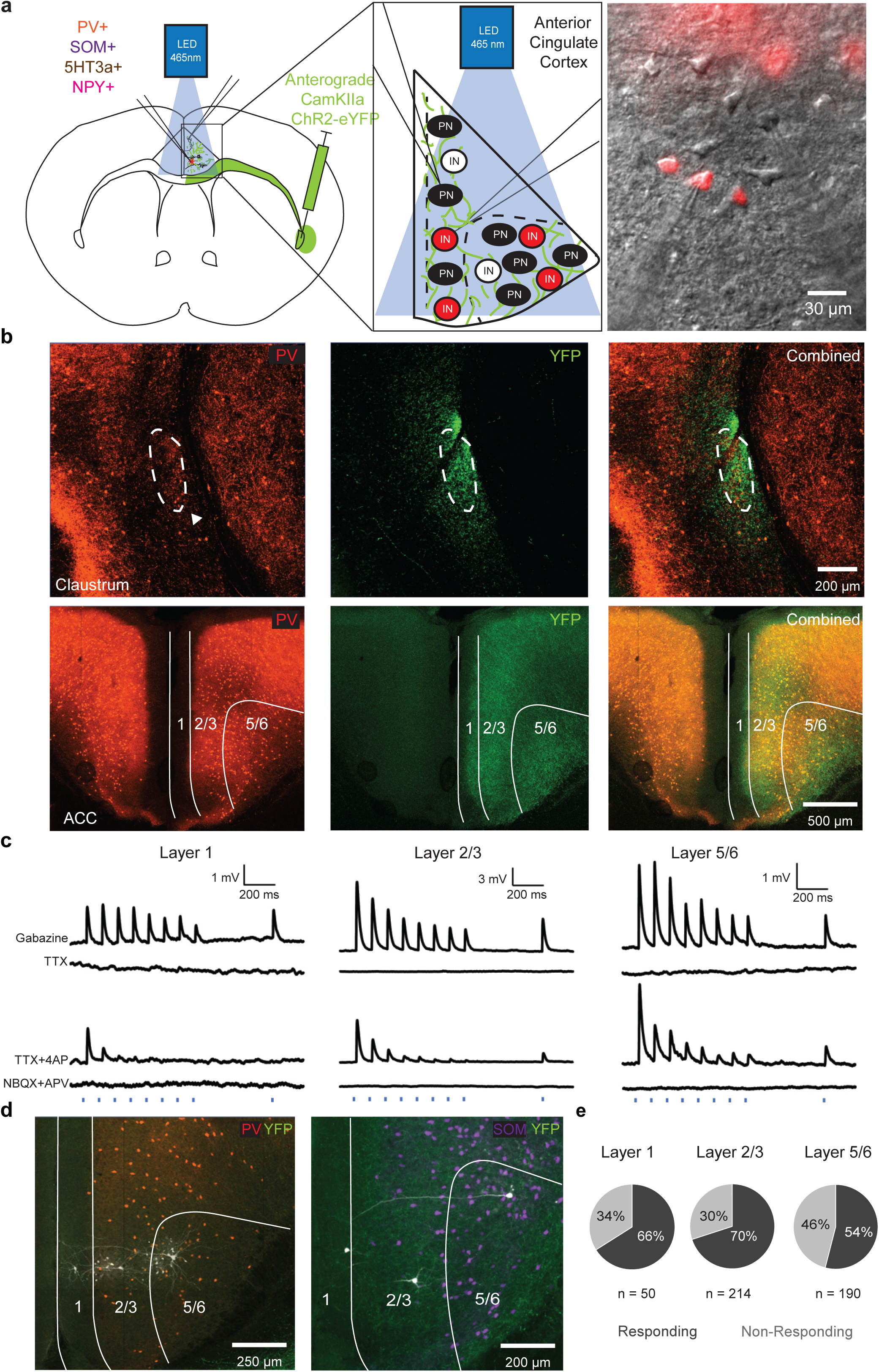
All layers of the ACC receive excitatory, monosynaptic input. **a**. Illustration depicting experimental paradigm. Briefly, ChR2 was injected using an anterograde viral vector into the claustrum (100nl, 300nl/min) into different transgenic mouse lines with florescent reporters to identify interneuron populations. Paired whole-cell patch clamp recordings were made in the ACC, and claustrum axons were photostimulated to investigate fore postsynaptic response. DIC image of paired recordings of interneuron and putative pyramidal neuron. **b**. Top: Confocal image of a 250 μm experimental slice, showing injection into claustrum with minor spillage into surrounding regions in a PV-tdTomato mouse. Dotted lines mark out the claustrum core identified by PV-rich neuropil staining. Bottom: Confocal image of a 250 μm experimental slice in a PV-tdTomato mouse, with a close-up in the ACC region showing YFP fibers in ipsilateral ACC but not contralateral ACC. **c**. Postsynaptic responses from layer 1, 2/3 and 5/6 neurons are recorded in Gabazine. The postsynaptic response is subsequently removed on bath application of TTX and recovered with TTX+4AP, this confirms that the input is monosynaptic. The subsequent postsynaptic response is abolished with AMPA and NMDA receptor antagonists, NBQX and APV, confirming that the input is excitatory. **d**. Confocal image of a 250 μm experimental slice in a PV-tdTomato mouse and SOM-tdTomato mouse, showing filled neurons in layers 1, 2/3 and 5/6 respectively. **e**. Combined responses probability from all cells recorded in layers 1 (33 responding cells out of 47 recordings), 2/3 (166/223 recordings) and 5/6 respectively (139/235 recordings), there was no significant different in response probability across layers (*P* > 0.05, chi-squared test)

Whole-cell patch clamp recordings were performed in neurons located in layers 1, 2/3 and 5/6 of the ipsilateral ACC (Fig. 3b). To prevent feedforward inhibitory activity^13^, GABA_A_ receptor blocker gabazine (10 μM) was included in the bath solution. A subset of responsive cells in all tested layers were probed for monosynaptic connections. In all layers tested, the postsynaptic response was abolished with Na+ channel blocker TTX (1 μm) and recovered with K+ channel blocker 4AP (100 μM), proving that monosynaptic connections are made from the claustrum to layer 1, 2/3 and 5/6 of the ACC (n = 5 recordings tested in layer 1; n = 20 recordings tested in layer 2/3, n = 27 recordings tested in layer 5/6; Fig. 3c, Supplementary Fig. 3). The same responding cells were then bath exposed to NMDA and AMPA receptor antagonists NBQX (10 μM) and APV (50 μM). Postsynaptic responses were abolished after bath application of NBQX and APV, confirming that the output from the CLA was indeed excitatory (n = 2 recordings in layer 1; n = 16 in layer 2/3, n = 8 in layer 5/6; Fig. 3c).

To verify experimenter layer attribution, a random selection of neurobiotin-filled ACC cells was stained with Cy5-dye and imaged (Fig. 3d). These images were compared post-hoc with the Allen Institute Reference Atlas^66^ and verified to be correctly assigned. All ACC layers received CLA input at different probabilities which were not due to chance (n = 33/47 recordings in layer 1; n = 166/223 in layer 2/3; n = 139/235 in layer 5/6; Fig. 3e, *P* < 0.05, Chi-squared test). Additionally, layer 6b neurons in the ACC (Supplementary Fig. 3, n = 3) were also observed to receive CLA input. In summary, the CLA sends excitatory, monosynaptic input to all layers of the ipsilateral ACC.

### Cell-type specificity in claustral projections to ACC

Three main populations of interneurons account for the majority of GABAergic neurons in the cortex – PV, SOM and 5HT3a expressing interneurons^59, 67^. PV interneurons facilitate feedforward inhibition^63^ and control of excitatory and inhibitory balance within the cortex^60, 61^; SOM interneurons receive facilitating excitation from cortical pyramidal neurons, target the dendrites of pyramidal neurons, and suppress the activity other interneurons^64^; 5HT3a neurons receive thalamocortical input and mainly inhibit superficial layers of the cortex^62^. A majority (69%) of neurons recorded from the PV population received CLA input (n = 52/75 recordings, Fig. 4a), 35% of neurons recorded from the SOM population received CLA input (n = 17/48, Fig. 4b), 58% of neurons recorded from the 5HT population received CLA input (n = 36/62, Fig. 4c). NPY interneurons are a unique population within the 5HT3a family^67^ that was previously suggested to play an important role in claustro-cortical feedforward inhibition^13^. 77% of neurons recorded from the NPY population received CLA input (n = 51/66, Fig. 4d).

**Fig. 4.**
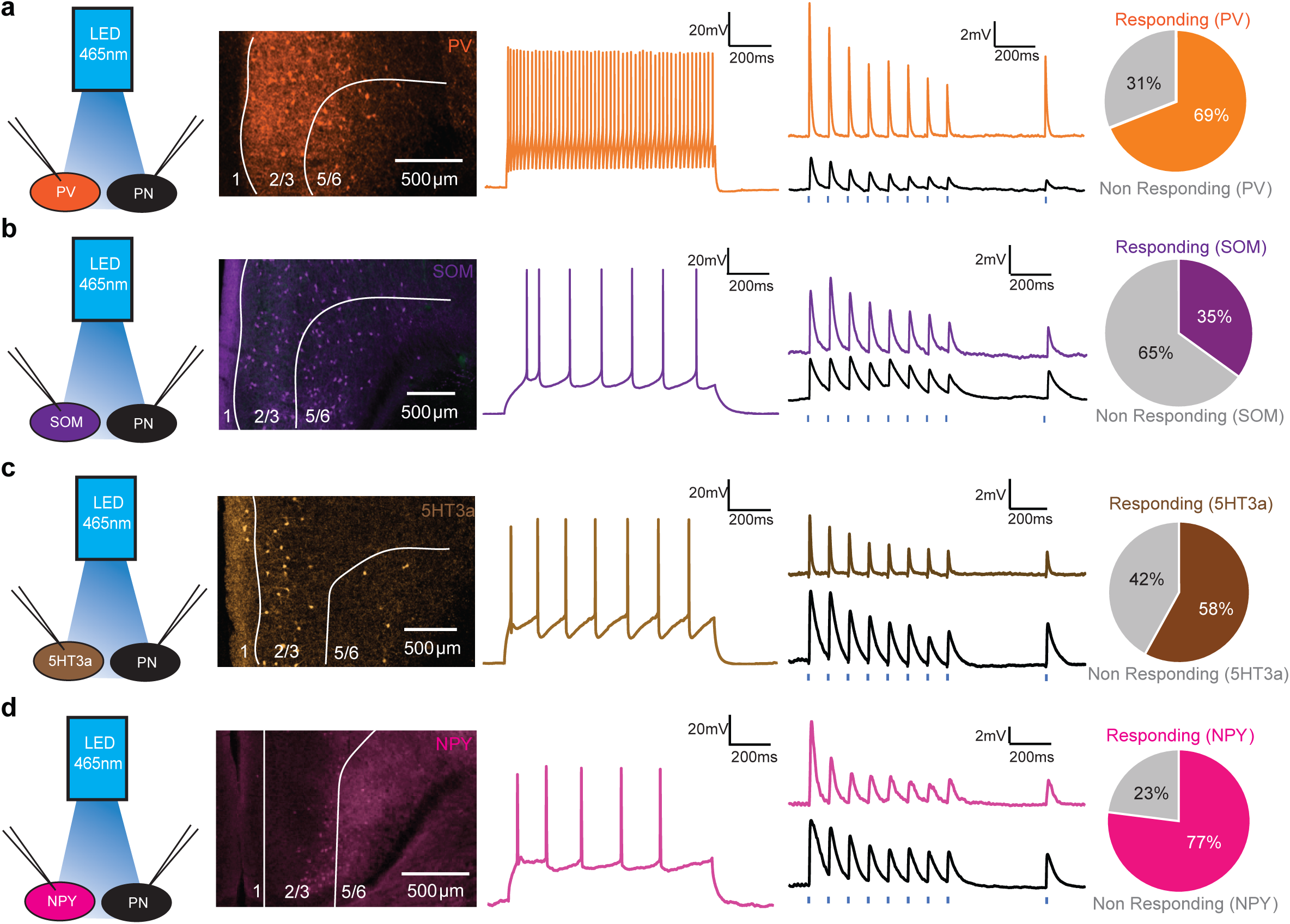
Claustrum projections selectively target interneuron populations. **a**. Illustration of paired recordings in of PV interneurons and pyramidal neurons. Confocal image of the ACC shows that PV interneurons are in layers 2/3 and 5/6 of the ACC, a total of 75 PV interneurons (42 in layer 2/3, 33 in layer 5/6) were studied. Example trace of PV interneuron. Representative postsynaptic responses in paired recordings between PV and pyramidal neuron populations (black trace). 69% of all recorded PV neurons were found to be responsive. **b**. SOM: Confocal image of the ACC shows that SOM interneurons are in layers 2/3 and 5/6 with neurites in layer 1, a total of 30 SOM interneurons (1 in layer 1, 14 in layer 2/3, 33 in layer 5/6) were studied. Example trace of SOM interneuron. Representative postsynaptic responses in paired recordings between SOM and pyramidal neuron populations (black trace). 35% of all recorded SOM neurons were found to be responsive. **c**. 5HT3a: Confocal image of the ACC shows that 5HT3a interneurons are located mostly in layers 1 and layers 2/3 with sparse labelling in layer 5/6. A total of 62 5HT3a interneurons (19 in layer 1, 27 in layer 2/3, 16 in layer 5/6) were studied. Example trace of 5HT3a interneuron. Representative postsynaptic responses in paired recordings between 5HT3a and pyramidal neuron populations (black trace). 58% of all recorded 5HT3a neurons were found to be responsive. **d**. NPY: Confocal image of the ACC shows that NPY interneurons are in layers 1 and layers 2/3 with heavy labelling in layer 5/6. A total of 66 NPY interneurons (10 in layer 1, 27 in layer 2/3, 29 in layer 5/6) were studied. Example trace of NPY interneuron. Representative postsynaptic responses between NPY and pyramidal neuron populations (black trace). 77% of all recorded NPY neurons were found to be responsive.

While PV neurons in both ACC layers 2/3 and 5/6 received CLA input evenly (n = 31/42 recordings, in layer 2/3; n = 21/33 in layer 5/6; *P* > 0.05, chi-squared test; Fig. 5a, b) there were differences in their postsynaptic responses. PV interneurons in layer 2/3 showed significantly shorter response latency compared to pyramidal neurons in the same layer (PV: 3.33 ± 0.16 ms, non-PV: 4.09 ± 0.21 ms, *P* < 0.001, t-test; PV: 3.62 ± 0.32 ms, non-PV: 3.95 ± 0.42 ms, *P* < 0.05, paired t-test; Fig. 5c) but had comparable response latencies in layer 5/6 (PV: 2.8 ± 0.15 ms, non-PV: 3.61 ± 0.17 ms, *P* < 0.01, t-test; PV: 2.94 ± 0.24 ms, non-PV: 3.52 ± 0.36 ms, *P* > 0.05, paired t-test; Fig. 5c). The EPSP amplitude in layer 2/3 PV interneurons was not significantly different from the EPSP amplitude of putative layer 2/3 pyramidal neurons (PV: 5.1 ± 0.88 mV, non-PV: 5.04 ± 0.89 mV, *P >*0.05, t-test; PV: 3.85 ± 0.95 mV, non-PV: 6.04 ± 2.08 mV, *P >* 0.05, paired t-test, Fig. 5D), on the other hand, PV interneurons in layer 5/6 had significantly larger EPSP sizes (PV: 9.12 ± 1.34 mV, non-PV: 2.81 ± 0.38 mV, *P <* 0.001, t-test; PV: 8.15 ± 1.9 mV, non-PV: 3.29 ± 0.86 mV, *P <* 0.05, paired t-test, Fig. 5D). In summary, while the CLA targets both superficial and deep layers of the ACC, the post-synaptic responses of the PV interneurons differ in each layer.

**Fig. 5.**
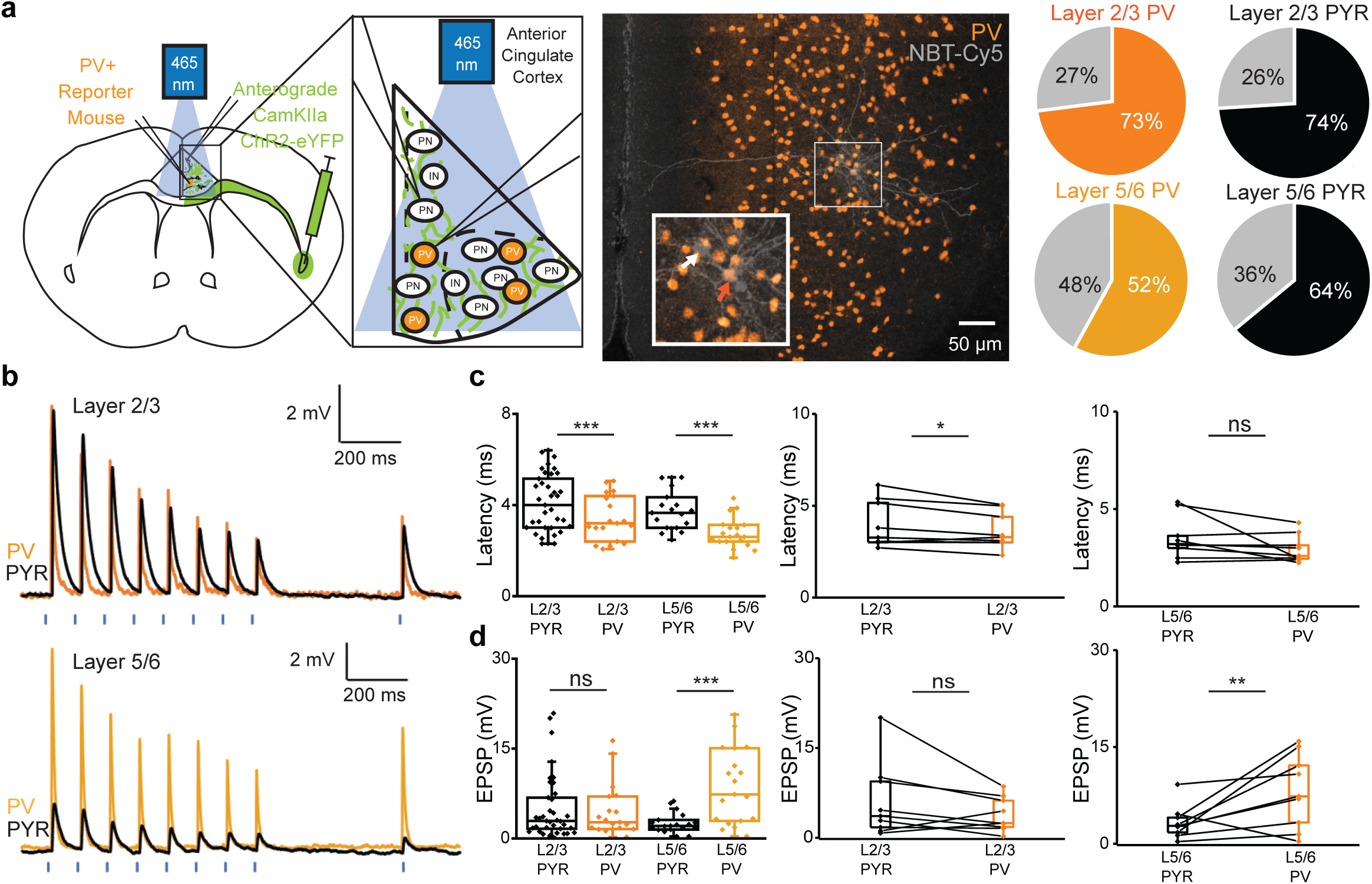
PV interneurons have layer-dependent postsynaptic responses. **a**. Illustration of experimental paradigm, briefly, paired whole-cell patch clamp recordings were made from neighboring PV interneurons and pyramidal neurons and subsequently photostimulated at 10 Hz. 74% (31/42 neurons) of PV neurons and 73% (41/56 neurons) of pyramidal neurons recorded in layer 2/3 responded to photostimulation. 64% (21/33 neurons) of PV neurons and 58% (33/57 neurons) of pyramidal neurons recorded in layer 5/6 responded to photostimulation. **b**. Representative EPSP traces of paired PV and pyramidal neurons to stimulation of claustrum projections (Dark orange: layer 2/3 PV; Light orange: layer 5/6 PV; Black: Pyramidal neurons) at 10Hz frequency. **c**. Layer 2/3 neurons have shorter latency than pyramidal neurons (*P* < 0.01, t-test) at the population level, a result confirmed in paired recordings (*P* < 0.05, paired t-test). Although Layer 5/6 neurons have shorter latency than pyramidal neurons (*P* < 0.01, t-test) at a population level, this result is not confirmed in paired recordings (*P* > 0.05, paired t-test). **d**. EPSP size was significantly different in layer 5/6 PV cells compared to pyramidal neurons (*P <* 0.05, t-test) but not in layer 2/3 (*P >* 0.05, t-test). This observation is further confirmed in paired recordings (*P >* 0.05 in layer 2/3, *P <* 0.05 in layer 5/6, paired t-test).

Although SOM neurons in both layers responded to photostimulation of CLA projections the difference in response rate was stark (n = 1/14 recordings in layer 2/3; n = 15/33 in layer 5/6, Fig. 6a,b), suggesting that CLA preferentially targets the layer 5/6 of the ACC. Paired recordings of SOM interneurons and pyramidal neurons were made in layer 5/6 of the ACC. SOM interneurons and pyramidal neurons shared comparable response latency (SOM: 2.89 ± 0.25 ms, non-SOM: 2.97 ± 0.22 ms, *P >* 0.05, t-test; SOM: 2.74 ± 0.41 ms, non-SOM: 2.88 ± 0.2 ms, *P >* 0.05, paired t-test, Fig. 6c). The EPSP amplitude in SOM interneurons was likewise, not significantly different from that observed in pyramidal neurons (SOM: 9.81 ± 2.31 mV, non-SOM: 6.11 ± 1.43 mV, *P >* 0.05, t-test; SOM: 11.06 ± 3.95 mV, non-SOM: 5.17 ± 2.35 mV, *P >* 0.05, paired t-test, Fig. 6d). Taken together our data suggest that the CLA preferentially targets SOM interneurons in the infragranular ACC although these SOM interneurons are similar in response latency and EPSP amplitude to their pyramidal neurons in layer 5/6.

**Fig. 6.**
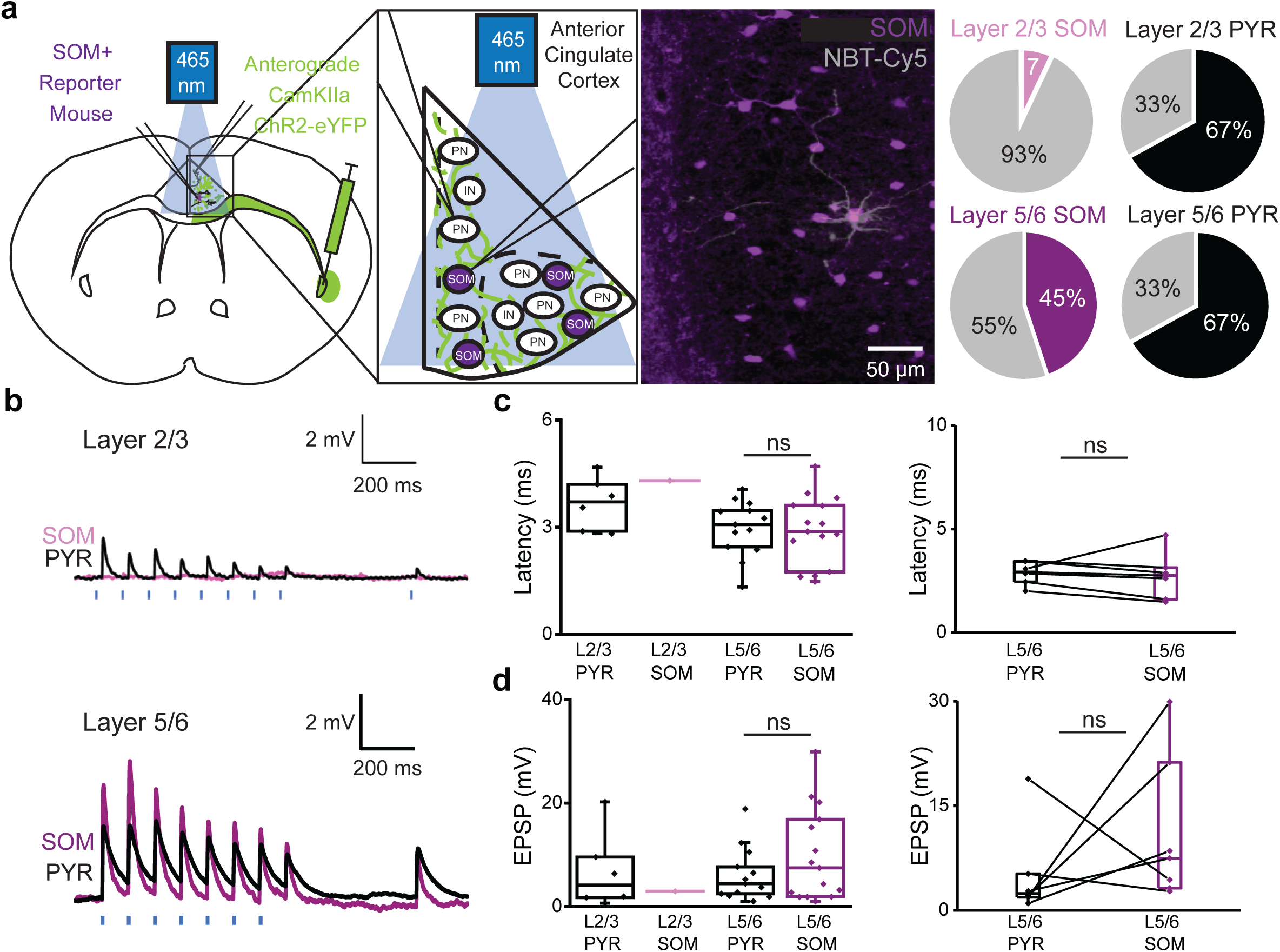
Claustrum projections mainly target infragranular SOM interneurons. **a**. Illustration of experimental paradigm in SOM+ reporter mice. 7% (1/14 neurons) of SOM neurons and 67% (8/12 neurons) of pyramidal neurons recorded in layer 2/3 responded to photostimulation, while 45% (15/33 neurons) of SOM neurons and 67% (14/21 neurons) of pyramidal neurons recorded in layer 5/6 responded to photostimulation. **b**. Representative EPSP traces of paired SOM and pyramidal neurons to stimulation of claustrum projections (Light purple: layer 2/3 SOM; Dark purple: layer 5/6 SOM; Black: Pyramidal neurons) at 10Hz frequency. **c**. No significant difference in response latency between cells in layer 5/6 (*P* > 0.05, t-test). This observation is further confirmed in paired recordings (*P* > 0.05, paired t-test). **d**. No significant difference in response latency between cells in layer 5/6 (*P* > 0.05, t-test). This observation is further confirmed in paired recordings (*P* > 0.05, paired t-test).

Both layer 2/3 and layer 5/6 5HT3a neurons likewise received CLA input although there was a marked bias in favor of layer 2/3 5HT3a neurons (n =20/27 in layer 2/3; n = 2/16 in layer 5/6; p < 0.001, chi-squared test, Fig. 7a,b). Layer 1 neurons expressing 5HT3a-receptor marker were also found to receive claustrum input (Supplementary Fig. S3). Paired recordings of 5HT3a interneurons and pyramidal neurons were only made in layer 2/3 of the ACC. Layer 2/3 5HT3a interneurons and pyramidal neurons responded to CLA input with the same latency (5HT3a: 3.42 ± 0.36 ms, non-5HT3a: 2.87 ± 0.14 ms, *P >* 0.05, t-test; 5HT3a: 2.5 ± 0.17 ms, non-5HT3a: 2.73 ± 0.19 ms, *P >* 0.05, paired t-test, Fig. 7c). The EPSP amplitude in layer 2/3 5HT3a interneurons was not significantly different from that observed in pyramidal neurons (5HT3a: 7.26 ± 2.04 mV, non-5HT3a: 5.66 ± 0.79 mV, *P >* 0.05, t-test; 5HT3a: 6.76 ± 1.35 mV, non-5HT3a: 5.84 ± 1.61 mV, *P >* 0.05, paired t-test, Fig. 7d). Our data supports the conclusion that CLA preferentially targets 5HT3a neurons in the superficial ACC but not the deep layer ACC, and that the response of these neurons is not significantly faster or stronger than the corresponding pyramidal neurons in the same layer.

**Fig. 7.**
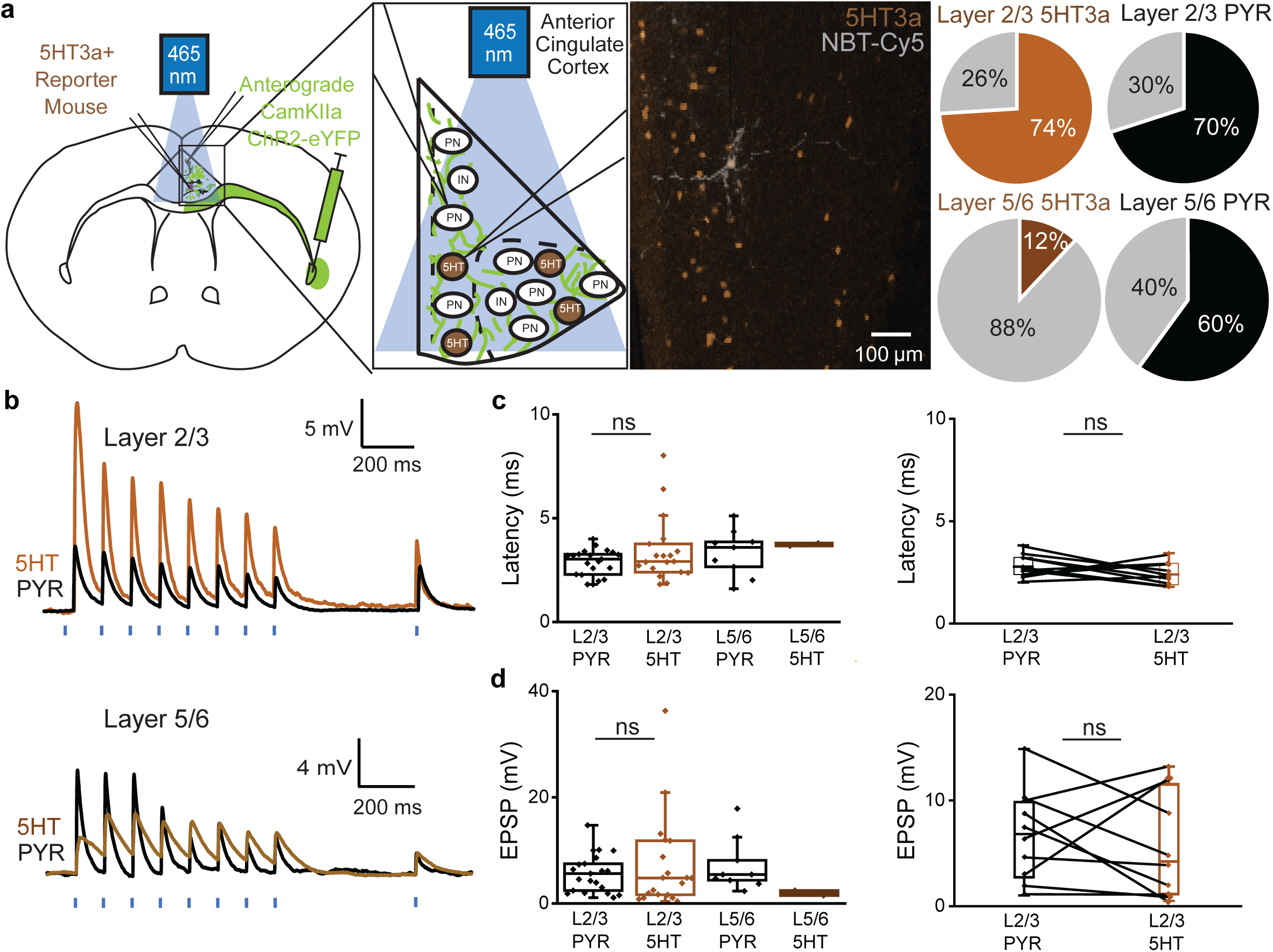
Claustrum projections mainly target supragranular 5HT interneurons. **a.** Illustration of experimental paradigm in 5HT3a+ reporter mice. 74% (20/27 neurons) of 5HT3a neurons and 71% (22/31 neurons) of pyramidal neurons recorded in layer 2/3 responded to photostimulation. 13% (2/16 neurons) of 5HT3a neurons and 60% (9/15 neurons) of pyramidal neurons recorded in layer 5/6 responded to photostimulation. **b**. Representative EPSP traces of 5HT3a and pyramidal neurons to stimulation of claustrum projections (Light brown: layer 2/3 5HT3a; Dark brown: layer 5/6 5HT3a; Black: Pyramidal neurons) at 10Hz frequency. **c.** No significant difference in response latency between cells in layer 2/3 (*P* > 0.05, t-test). This observation is further confirmed in paired recordings (*P* > 0.05, paired t-test). **d.** No significant difference in EPSP amplitude between cells in layer 2/3 (*P* > 0.05, t-test). This observation is further confirmed in paired recordings (*P* > 0.05, paired t-test).

A very high percentage of the NPY neurons responded to photostimulation of CLA projections in both layer 2/3 and 5/6 (n = 19/27 in layer 2/3; n = 25/29 in layer 5/6; Fig. 8,b). Layer 1 neurons expressing NPY marker were also found to receive claustrum input (Supplementary Fig. 3). NPY interneurons and pyramidal neurons in the same layer responded to CLA input with comparable latency in both Layers 2/3 (NPY: 3.39 ± 0.54 ms, non-NPY: 3.92 ± 0.52 ms, *P >* 0.05, t-test; NPY: 3.58 ± 0.82 ms, non-NPY: 4.1 ± 0.97 ms, *P >* 0.05, paired t-test, Fig. 7c) and layers 5/6 (NPY: 3.47 ± 0.28 ms, non-NPY: 3.19 ± 0.13 ms, *P >* 0.05, t-test; NPY: 3.41 ± 0.47 ms, non-NPY: 3.14 ± 0.18 ms, *P >* 0.05, paired t-test, Fig. 7c). The EPSP amplitude in layer 2/3 NPY interneurons was not significantly different from that observed in pyramidal neurons (NPY: 8.85 ± 1.83 mV, non-NPY: 7.47 ± 1.26 mV, *P >* 0.05, t-test; NPY: 10.66 ± 2.51 mV, non-NPY: 8.62 ± 2.1 mV, *P >* 0.05, paired t-test, Fig. 8d), this was also the case for layer 5/6 NPY-pyramidal comparison (NPY: 8.23 ± 1.1 mV, non-NPY: 5.81 ± 0.93 mV, *P >* 0.05, t-test; NPY: 10.32 ± 1.39 mV, non-NPY: 7.36 ± 1.08 mV, *P >* 0.05, paired t-test, Fig. 8d). In conclusion, the CLA provides direct monosynaptic input to all tested neuron types in the ACC, however, the probability of connectivity and relative amplitudes depended on both the postsynaptic neuron type and the cortical layer.

**Fig. 8.**
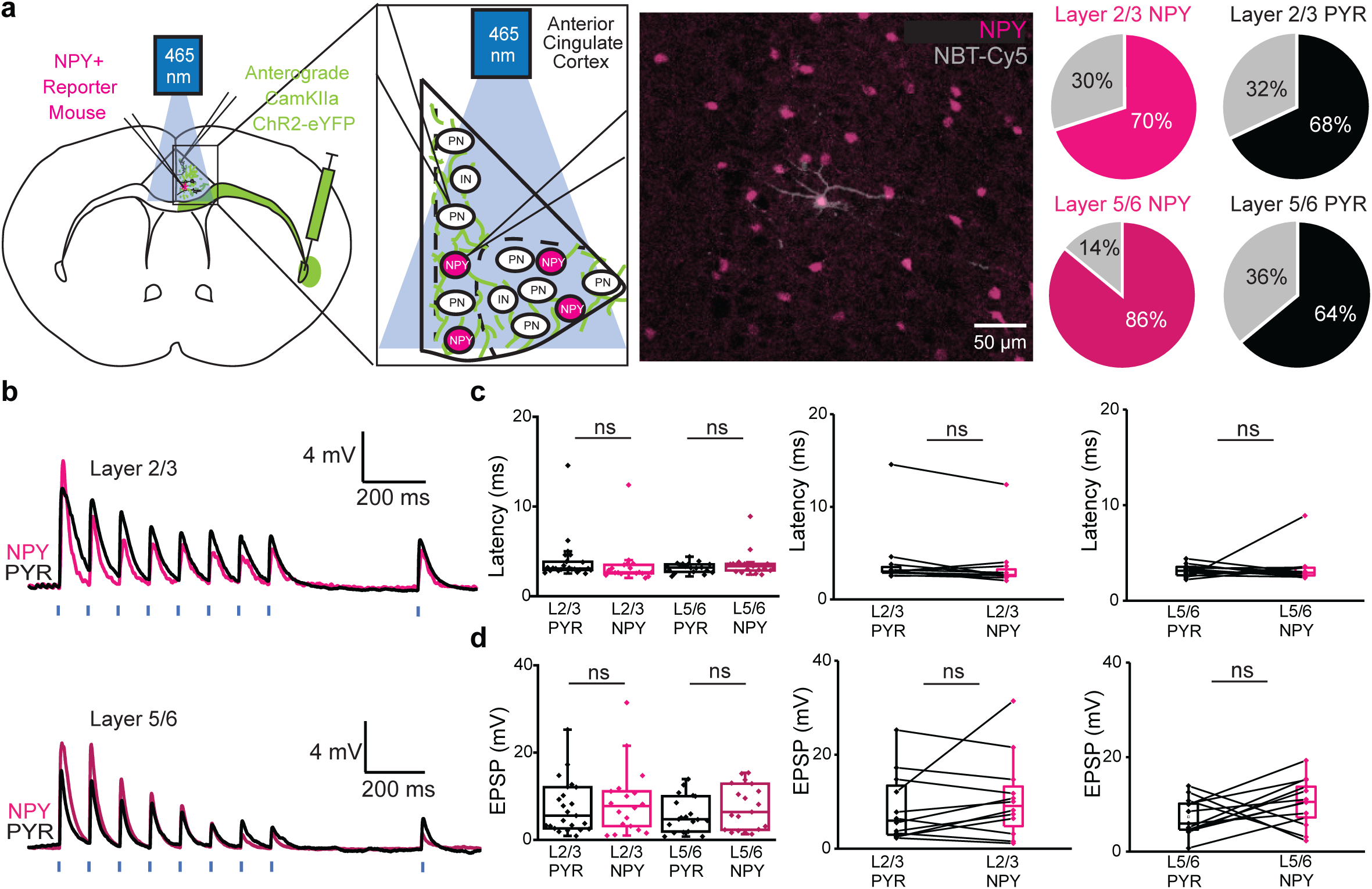
Supragranular and infragranular NPY interneurons do not response faster or larger to claustrum input. **a.** Illustration of experimental paradigm in NPY+ reporter mice. 70% (19/27 neurons) of NPY neurons and 69% (24/35 neurons) of pyramidal neurons recorded in layer 2/3 responded to photostimulation. 65% (20/31 neurons) of NPY neurons and 60% (9/15 neurons) of pyramidal neurons recorded in layer 5/6 responded to photostimulation. **b.** Representative EPSP traces of NPY and pyramidal neurons to stimulation of claustrum projections (Light pink: layer 2/3 NPY; Dark pink: layer 5/6 NPY; Black: Pyramidal neurons) at 10Hz frequency. **c.** No significant difference in response latency within layers (*P* > 0.05, t-test). This observation is further confirmed in paired recordings (*P >* 0.05 in both situations, paired t-test). **d.** No significant difference in EPSP amplitude within layers (*P* > 0.05, t-test). This observation is further confirmed in paired recordings (*P >* 0.05 in both situations, paired t-test).

### VGLUT2-expressing claustrum projection neurons selectively target supragranular ACC

Recent studies suggest VGLUT2 to be a selective marker for CLA projection neurons^45, 68^. To investigate how this specific CLA projection population targets the ACC, Cre-dependent ChR2 was injected into the CLA region and red retrograde florescent beads were injected into the ACC of VGLUT2-Cre transgenic mice. *Ex vivo* whole-cell patch clamp recordings were subsequently performed in the ACC (Fig. 9a). Florescent ChR2 fibers were observed in all layers of the ipsilateral ACC. Red retrobeads injected in the ACC were observed in VGLUT2 expressing CLA neurons (RetroBead+/VGLUT2+) as well as neurons that did not express VGLUT2 (RetroBead+/VGLUT2-, Fig. 9a), suggesting that the CLA-ACC pathway is mediated by both VGLUT2 positive and negative projection neurons.

**Fig. 9.**
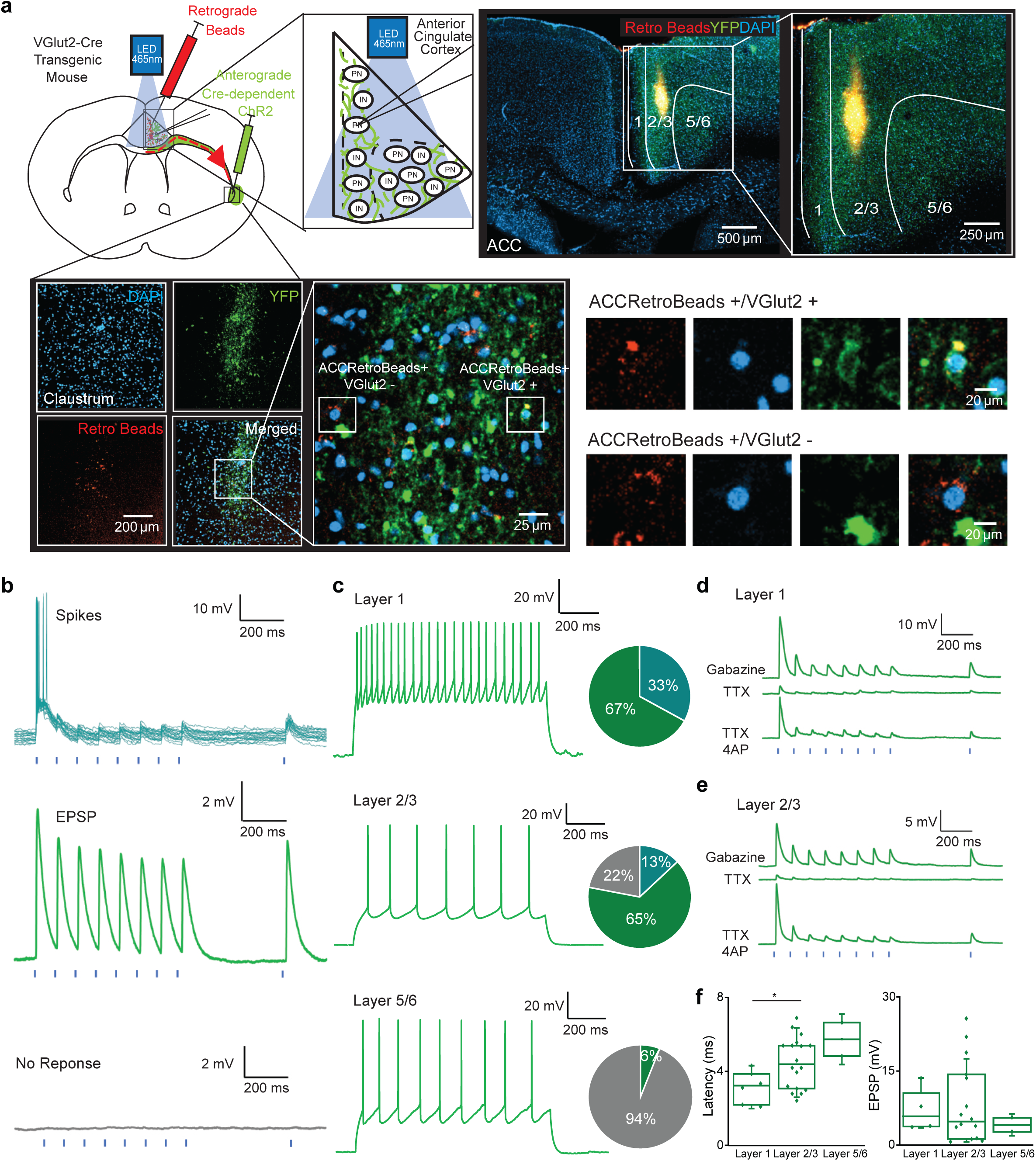
VGLUT2-expressing CLA neurons selectively target superficial ACC layers. **a.** Top left: Illustration of experimental paradigm, briefly, retrograde fluorescent beads were injected into the ACC, and Cre-dependent ChR2 was injected into the claustrum of VGLUT2-Cre transgenic mice. Whole-cell patch clamp recordings were made in all layers of the ACC and subsequently photostimulated at 10 Hz. Top right: Confocal image of a 250 μm experimental slice, showing injection of red retrograde beads into the ACC, and claustrum axons at the ACC. Bottom left: Confocal image of the same 250 μm experimental slice, showing VGLUT2 expression and CLA-ACC neurons in the claustrum. Bottom right: two types of CLA-ACC neurons were observed (VGLUT2+ and VGLUT2-). **b**. Three types of response to photostimulation of CLA afferents: spikes, EPSP and no-response. **c**. Example trace of neurons recorded at layers 1, 2/3 and 5/6 respectively. 100% (6/6 neurons) of neurons recorded in layer 1 responded to photostimulation, 78% (18/23 neurons) of neurons recorded in layer 2/3 responded to photostimulation, while 6% (2/33 neurons) out of neurons in layer 5/6 responded to photostimulation, this difference in distribution was not due to chance (*P <* 0.001, chi-squared test). **d.** A subset of layer 1 neurons (n =3) and **e.** layer 2/3 (n= 11) that responded to photostimulation were tested for monosynaptic connections, postsynaptic response to photostimulation was abolished with bath application of TTX and recovered upon 4AP application. **f.** Layer 1 neurons responded with a significantly shorter latency compared to layer 2/3 neurons (*P <* 0.05, t-test).

All neurons recorded in layer 1 responded to photostimulation (n = 6/6 recordings), two of which depolarized beyond spiking threshold in response to photostimulation of the same intensity (Fig. 9b-d). Most of the recorded neurons in Layer 2/3 responded to photostimulation (78%, n = 18/23, Fig. 9c), 4 of which depolarized beyond spiking threshold. In contrast, only a small fraction on neurons in Layer 5/6 responded to photostimulation (6%, n = 2/31 recordings, Fig 9c). The rate of VGLUT2 input is, therefore, significantly higher for layer 2/3 rather than layers 5/6 (*P <* 0.001, One Way ANOVA). In a subset of responding neurons the postsynaptic response was abolished with TTX (1 μm) and recovered with K+ channel blocker 4AP (100 μM), indicating that monosynaptic connections are formed by VGLUT2 expressing neurons onto the supragranular ACC (n=3 in layer 1, Fig. 9d; n=11 in layer 2/3, Fig. 9e). The response latency of layer 1 recordings was significantly shorter than those in layer 2/3 (p < 0.05, t-test, Fig. 9f). Statistical tests were not performed for Layer 5/6 responses due to the low number of responding neurons. In these experiments interneurons did not express fluorescent markers, therefore, VGLUT2recorded neurons were all putative pyramidal neurons. Taken together, our data suggests that VGLUT2-expressing CLA projection neurons selectively target supragranular layers of the ACC in comparison to the broad innervation provided by CaMKIIa-expressing CLA-ACC projections.

**Fig. 10.**
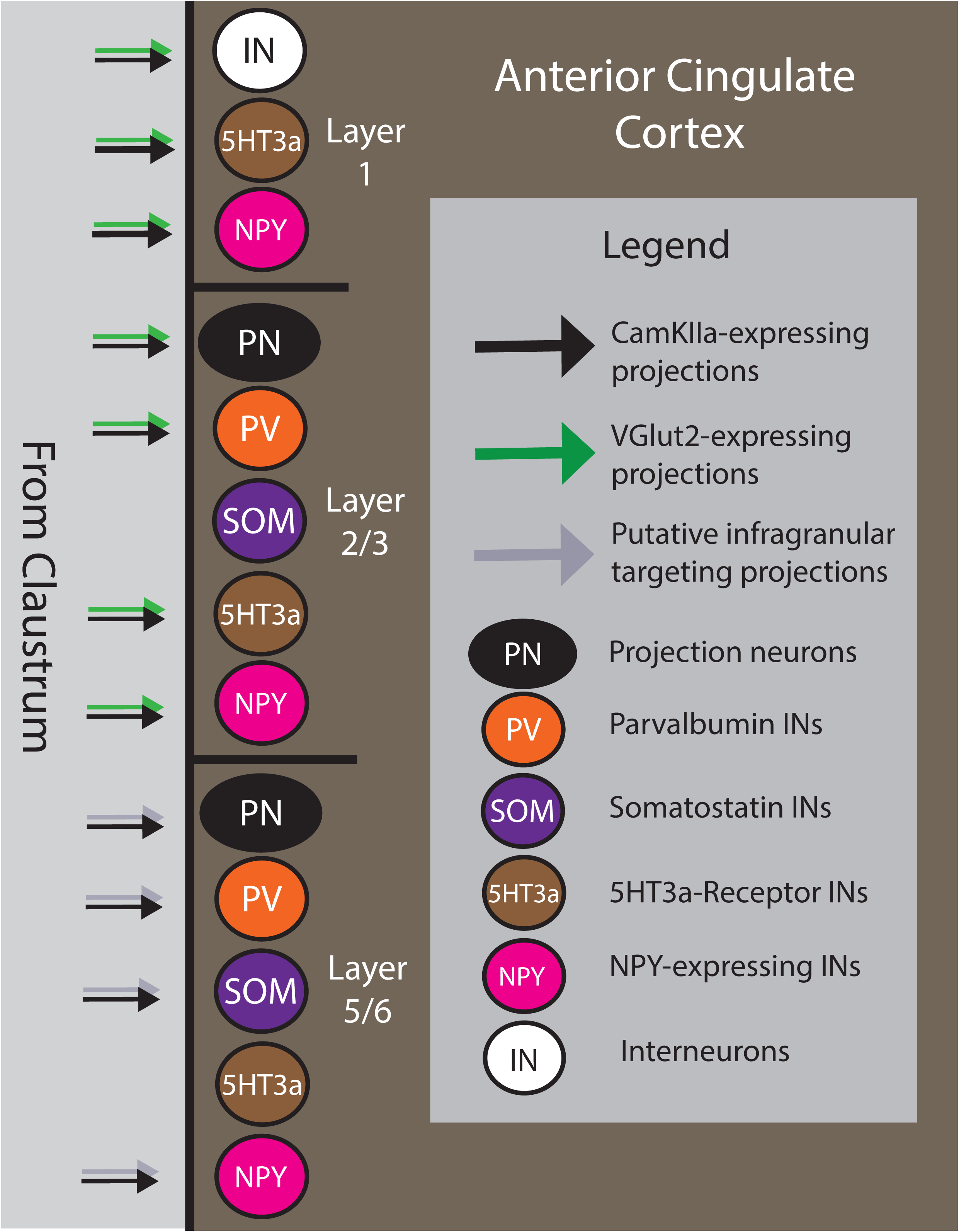
Summary scheme: claustrum projections to the ACC are layer and cell-type dependent. Illustration showing different claustrum targets in the ACC. Claustrum projections were observed to target all cell types in layer 1. PV, 5HT3a and NPY neurons in layer 2/3 as well as PV, SOM and NPY neurons in layer 5/6. VGLUT2-expressing claustrum neurons targeted all neurons recorded in layer 1 and layer 2/3. A putative infragranular targeting claustrum projection neuron population may also exist.

## Discussion

In this study, we used a combination of *in vivo* and *ex vivo* electrophysiological methods to investigate the organizing principles, targets and influence of CLA projections to the ACC. We show that *in vivo* activation of CLA neurons produces a combination of excitation and inhibition that depends on three factors; CLA projection neuronal identity, ACC layer that receives the CLA input, and the cell type in the ACC. Using *ex vivo* experiments, we obtained a detailed mapping of the pathway revealing that excitatory monosynaptic CLA projections to ACC targeted PV interneurons in both layers 2/3 and 5/6, while preferentially targeting 5HT3a interneurons in the supragranular layers and SOM interneurons in the infragranular layers. The intricate synaptic organization of the CLA-ACC pathway may explain the complex dynamics seen in *in vivo* in the ACC following activation of of CLA.

### Modulation of ACC by broad activation of CLA neurons in vivo

CLA has been reported to exert either excitatory or an inhibitory effect on different cortical regions^1, 18^. There is anatomical and functional evidence showing that CLA outputs target both excitatory and inhibitory cells within cortical circuits^2, 10, 13^. In addition, the fact that CLA is composed by different neuron types, indicates that simultaneous activation of all projection neurons may account for the discrepancy observed regarding the influence of CLA on cortical processing^11, 12, 19^.

Our *in vivo* studies show broad activation with 5 ms light pulses of CLA neurons expressing ChR2 driven by the CaMKIIa promoter produce, in the LFP band, an overall small and brief excitation follow by a potent inhibition in the ACC. This type of response was observed both in layer 2/3 and layer 5/6 and suggested, as previously reported^13, 18, 50^ that CLA provides a “blanket of inhibition” to the ACC. However, LFP responses while easy to record, have proven notoriously difficult to interpret^69–71^. Analysis of unit recordings showed that CLA photostimulation triggered a combination of excitation and inhibition in the ACC. In this context, although FS units were more likely to be excited than RS ones, we also observed excitation of RS units mainly in deep layers (Fig 2g,h,j). On the other hand, the proportion of both RS and FS inhibited after photostimulation of CLA neurons was much larger than the excited ones, especially in the supragranular layer (Fig 2j). This inhibition is weaker than the near-total and prolonged silence of cortical neurons reported previously^13, 50^. This could be due to the different effects of CLA input on different cortical areas. A recent study showed that activation of Esr2 population, a subgroup of CLA neurons driven by the CaMKIIa promoter, showed moderate inhibitory effect albeit their preference for FS and little impact in RS^18^. This raises the possibility that Esr2-expressing neurons may express their own target selectivity in cortex.

Recent studies have shown that activation of certain populations of CLA neurons, e.g. VGLUT2, might exert net excitation in cortex in contrast to the previously reported blanket of inhibition^45^. We decided to explore the influence *in vivo* in the ACC after activation of the VGLUT2 CLA neuronal subpopulation. In our study, we observed similar proportions of excitation and inhibition in layer 2/3 when comparing photostimulation in CaMKIIa and VGLUT2 mice. However, when comparing layer 5/6, we observed significant differences. Brief activation of VGLUT2 CLA neurons resulted in the activation of a high proportion of FS neurons (45.45%) and few regular-spiking (RS) neurons (8.77%). The strong recruitment of FS neurons in this layer also led to a strong inhibition of intralaminar RS neurons, which was stronger than in CaMKIIa experiments. The observed differences in the activation of the ACC in layer 5/6 after photostimulation of VGLUT2 and the broad activation of CLA neurons may lead to conflicting conclusions. One possibility is that the relatively low number of FS neurons in both groups might account for these differences. However, if this were the case, the impact on the RS neurons in layer 5/6 should be similar between the two groups but our results showed a strong inhibition of RS5/6 in the VGLUT2 group, which suggests a different mechanism of action. It is important to note that while RS neurons are primarily composed of putative pyramidal neurons, they may also include various subtypes of interneurons.

### Selective targeting of ACC interneurons by CLA projections

To gain a comprehensive and accurate understanding of the differences between ACC responses to activation of CaMKIIa or VGLUT2 CLA neurons, we conducted *ex vivo* experiments enabling selection of specific neuronal populations in ACC. Our results show that CaMKIIa-expressing CLA projection neurons preferentially targeted PV interneurons in both layers 2/3 and 5/6, 5HT3a interneurons in the supragranular layers, and SOM interneurons in the infragranular layers. Additionally, they also targeted NPY interneurons, a unique population within the 5HT3a expressing interneurons^67^ that was previously suggested to play a crucial role in claustro-cortical feedforward inhibition^13^. It is possible that the activation of certain interneurons may result in the downstream recruitment of other interneuron population, which could potentially decrease inhibition through disynaptic or polysynaptic effects. It has been shown that NPY interneurons in the PFC activated by CLA neurons provide feedforward inhibition to pyramidal neurons and PV interneurons^13^. This would explain a decrease on activity (either units excited or inhibited) after broad activation of CLA neurons.

Our *ex vivo* data shows, in agreement with anatomical data reported in recent studies^45^, that VGLUT2-expressing CLA projection neurons preferentially targeted layer 2/3 of the ACC. With only two EPSP responses observed in layer 5/6 after CLA activation. Due to technical limitations with the VGLUT2-Cre mice, the reporter could not be expressed in any of the interneuron populations, hence all the neurons recorded in the *ex vivo* experiments were classified as putative pyramidal neurons. It is possible that the VGLUT2-expressing CLA population might target PV interneurons in layer 5/6, leading to the results observed *in vivo*. Another possibility to explain the results observed *in vivo* is that the excitation of the units may not be a direct effect of CLA activation, but instead could be a result of translaminar polysynaptic inputs ^72, 73^. Untangling the effect of VGLUT2-CLA populations on interneurons in an *ex vivo* setting would be relevant future work to testing these hypotheses.

In addition to the selective targeting of interneurons in layers 2/3 vs 5/6, we also show that 5HT3a and NPY expressing Layer 1 ACC interneurons also received CLA input. Layer 1 receives input from brain regions associated with higher cognitive functions, cholinergic and monoaminergic systems as well as the secondary sensory ascending system involved with attention and saliency processes^74–80^. These inputs modulate the activity of the cortical column ^81^ via complex local inhibitory and disinhibitory circuits^77, 79^. This supports recent findings that implicate CLA projections to attention and saliency processes^11, 43, 51^.

### Strengths and limitations of our experimental approach

We combined *in vivo* and *ex vivo* recordings in this study to map the functional impact of CLA on the ACC. The objective of the *in vivo* recordings was to analyze the net response of the ACC after receiving CLA input, whereas the objective of *ex vivo* recordings was to identify specific cellular populations that received these inputs. While *in vivo* experiments give a more physiological readout, the reader should note that awake *in vivo* recordings are strongly affected by ongoing brain activity and numerous parallel input pathways. We hence used a paired, whole-cell *ex vivo* approach to complement the *in vivo* results and characterize specific cellular input.

In this study we used fluorescent markers for several GABAergic interneuron populations, which were simultaneously recorded with neighboring pyramidal cells, however, cortical pyramidal cells are made up of different subpopulations^82^ that were not classified here. Our approach allowed the *ex vivo* data to be controlled for variations in experimental animal, viral titer, incubation time, viral expression, cell-type, and brain slice quality across and within experiments. To identify the various molecularly defined mouse lines, we crossed a Cre-dependent line with a tdTomato reporter. While this is a commonly used method, care should be taken to check if the molecularly defined marker is developmentally regulated or constitutively expressed to provide a clearer picture of the findings. Moreover, this targeting of classification of interneurons was possible only in the CaMKIIa experiments but not when using VGLUT2-Cre mice. Future studies are needed to establish the detailed targeting of cortical neurons by the different CLA projection types such as those expressing VGLUT2, VGLUT1, Esr2, and others. In this paper, we built on our previous work^10, 11^ by using the robust CLA-ACC pathway to investigate cell-type and layer targets of CLA input at the cortex. As there is growing understanding that the cortical microcircuits of these regions differ^83, 84^, similar work should be performed on other granular and agranular cortices before a canonical claustro-cortical scheme can be developed.

## Conclusion

We report that the CLA activation evokes synaptic responses in all layers of the ACC and that these responses depend on the identity of the presynaptic CLA projection, the identity of the postsynaptic neuron, and the targeted cortical layer. Our results explain the discrepancies regarding the excitatory or inhibitory influence of claustrum in ACC and lay the foundation for understanding the organizing principles of claustro-cortical connectivity.

## Biographical Note

Z.C was a PhD candidate in the KI-NTU Joint PhD Program. R.T.M is a postdoctoral researcher at KI; J.F.N and A.T are PhD candidates at the Department of Neuroscience, Karolinska Institutet, Sweden G.J.A is the Irene Tan Liang Kheng Professor of Neuroscience at the Lee Kong Chian School of Medicine, Singapore. G.S is Professor of Neurophysiology at the Karolinska Institutet, Sweden.

## Author contributions

Z.C organized and performed *ex vivo* experimental work and data analysis. R.T.M organized and performed *in vivo* experimental work and data analysis. A.T. performed and analyzed *ex vivo* experiments. J.F.N. analyzed *in vivo* data. Z.C, R.T.M, G.J.A and G.S. were responsible for experimental design. All authors contributed to manuscript preparation.

## Acknowledgements

We thank the Silberberg lab for helpful discussions. We also thank Ainara and Erik for all their support. This work was founded by a Wallenberg Fellowship from the Knut and Alice Wallenberg Foundation, ERC Starting grant Swedish Brain Foundation (Hjärnfonden) and Swedish Medical Research Council (VR-M) to G.S; a Singapore Ministry of Education [MOE2015-T2-2-095] grant to G.J.A.; R.d.l.T.M was supported by a Karolinska Institute postdoctoral scholarship and StratNeuro postdoctoral grant; Z.C. was supported by the Nanyang Technological University - Karolinska Institutet Joint PhD Programme; A.T. and J.F.N were supported by a Karolinska Institute doctoral grant.

## Figure Legends

Supplementary Fig. 1 (related to Fig. 1 and 2) Optogenetic identifiication and stimulation of VGLUT2 CLA neurons in awake head-restrained mice.

Schematic of the in vivo recordings and photostimulation of the VGLUT2 neurons in the CLA. Raster plot showing the robust response (50 repetitions) of VGLUT2-CLA units in response to 5 ms and 500 ms light stimulation.

Supplementary Fig. 2 (related to Fig. 1) ChR2+ expression outside of the CLA did not result in any functional responses or photoelectrical artifacts in the ACC following CLA photostimulation in awake head-restrained mice.

Confocal image of a 250 μm experimental slice in a PV-tdTomato mouse, with a close-up in the CLA (left panel) and ACC region (right panel) showing virus injection outside the CLA and lack of YFP expression in the fibers in the ipsilateral ACC. In red, the optic fiber (left panel) and the silicon probe (right panel) are indicated. Heat map showing 150 repetitions with no LFP responses in the ACC layer 2/3 to 5 ms photostimulation of CLA neurons driven by a CaMKIIa promoter.

Supplementary Fig. 3. Layers 1 and 6b neurons in the ACC also receive claustrum input.

**a**. Example of neurobiotin-filled layer 6b neuron. **b**. Layer 6b neurons in the ACC received monosynaptic input from the claustrum (n = 3). **c**. Representative EPSP traces of undefined, SOM and 5HT3a layer 1 interneurons to stimulation of claustrum projections at 10Hz photostimulation rate. **d**. Percentage of Layer 1 recordings that respond to claustrum input: undefined PSNs (50%, 8/16 recordings) SOM (100% 1/1 recording), 5HT3a (66%, 8/12 recordings), NPY (20%, 1/5 recordings). **e**. No significant difference in response latency between cells in layer 1 (*P >* 0.05, One Way ANOVA). **f**. No significant difference in EPSP amplitude between cells in layer 1 (*P >* 0.05, One Way ANOVA). **g**. Postsynaptic responses from different interneuron types are recorded in gabazine. The postsynaptic response is subsequently removed on bath application of TTX and recovered with TTX+4AP, this confirms that the input is monosynaptic.

## Materials and Methods

### Animals

All experiments were performed according to the Guidelines of the Stockholm municipal committee for animal experiments. 26 adult PV-Cre ^85^, 5 adult SOM-Cre mice ^86^, 12 adult 5HT3a-Cre ^87^ mice and 15 adult NPY-Cre ^88^ were crossed with tdTomato reporter mouse line^89^ to obtain transgenic interneuron reporter mice. Another 4 adult NPY-Cre mice were used to compare differences between NPY-Cre and NPY-tdTomato mouse lines. 6 adult VGLUT2-Cre^90^ transgenic mice of both sexes were used to study the effect of VGLUT2-expressing claustrum projection neurons on the ACC. Mice were at least one month old before surgery was performed.

### Virus injection

Mice of either sex, 6-8 weeks old, were anesthetized with isoflurane and placed in a stereotaxic frame (Harvard Apparatus). To target the claustrum, a craniotomy was made −0.7 mm anterior-posterior (A/P) to bregma, + 3.4 mm medio-lateral (M/L) to the midline and 4.2 mm dorso-ventral (D/V) from the skull. A total of 100 nl of virus (AAV2-CaMKIIa-hChR2(H134R)-EYFP, Addgene 26969 for PV, SOM, 5HT and NPY transgenic mice; AAV5.EF1.dflox.hChR2(H134R)-eYFP.WPRE.hGH, Addgene 20298 for VGLUT2-Cre mice; Penn Vector Core, Philadelphia, PA, USA) was injected by a micropipette using Quintessential Stereotaxic Injector (Stoelting Europe, Dublin, Ireland) at a rate of 300 nl/min. To target the ACC a craniotomy was made 0.7 mm anterior to bregma, 0.25 mm lateral to midline and 1.4 mm ventral. Two types of injections were made in the ACC. To identify ACC-projecting claustrum neurons in the VGLUT2-Cre mice, red retrograde beads (Lumafluor) were injected into the ACC at the above-mentioned craniotomy; to identify mature PV, SOM, 5HT and NPY interneurons in the ACC 200 nl of virus (pAAV-hSyn-DIO-mCherry, Addgene 50459) was injected into the ACC. The pipette was held in place for 3 mins after the injection before being slowly retracted from the brain. Post injection analgesics were given (0.03 mg/kg buprenorphine).

### Head implants

Adult mice (2 to 3 months old) were anesthetized with isofluorane, and the head was fixed in a stereotaxic apparatus (Stoelting)^91^. Temgesic (0.1 mg/Kg) was administered before the surgery. The body temperature was maintained at 36 and 37°C by a heating pad. To avoid ocular dehydration, an ocular ointment (Viscotears 2 mg/g, Alcon) was applied over the eyes. To reduce any possible pain sensation, lidocaine was applied on the skin surface before the incision. After removing the skin covering the regions of interest, the bone was gently cleaned. Targeted regions for optic fiber placement and electrophysiological recordings were marked using stereotaxic coordinates on the surface of the skull. ACC craniotomy coordinates for recordings: ACC layer 2/3: +0.5 mm A/P and +0.2 mm M/L; ACC layer 5/6: +0.5 mm A/P and +0.5 mm M/L. Optic fiber craniotomy coordinates: +0.5 mm A/P and +0.2 mm M/L. Then, a thin layer of light-curing adhesive (Ivoclar Vivadent) was applied on the exposed skull. An aluminum metal head-post was fixed with dental cement Tetric Evo (Ivoclar Vivadent) between hemispheres close to mid-sagittal suture. A wall of dental cement was built along the edge of the bone and covering the implant. After the surgery, the animals were returned to their home cage.

### In vivo electrophysiology

Following a recovery period of at least three days after implantation, mice were habituated to being head-restrained over a period of 3-4 days. The recordings were performed up to 7 days after the habituation. On the day of the experiments, mice were anesthetized with isoflurane and small craniotomies (300-500 μm in diameter) were drilled to access the targeted areas. The open craniotomies were covered with silicone sealant (Kwik-Cast, WPI), and the animals were returned to their home cages for recovery. After 2 to 4 hours of recovery, mice were head-fixed, and the silicone from the craniotomies was removed. Extracellular LFP and spikes in the ACC were recorded using a silicon probe (A1×32-Poly2-5mm-50 s-177 and A1xOA32LP-Poly2-5mm-50 s-177, NeuroNexus, MI, USA) with 32 electrodes inserted 1.3 mm deep from the surface at the ACC L2/3 coordinates (+0.5 mm A-P; 0.25 mm M-L) or 1.5 mm deep from the surface at the ACC L5/6 coordinates (+0.5 mm A-P; 0.45 mm M-L). The probe was lowered using a micromanipulator (Luigs and Neumann, Germany) at a low speed (1 μm/s), to minimize damage to the brain tissue. To allow the brain to recover from the mechanical strain of the probe insertion, the recordings started 30 minutes after the probe was in its required position. The craniotomies were always covered with saline to avoid the dryness of the surface. The electrode reference was then connected to a silver wire positioned over the pia in a second craniotomy near the cerebellum with a second micromanipulator. The probe and the optic fiber were coated with DiI (1,1’-dioctadecyl-3,3,3′3’-tetramethylindocarbocyanine perchlorate, Invitrogen, USA) for post hoc recovery of the recording location. Recordings were performed using the OpenEphys systems. Broadband activity was sampled at 30 kHz (band pass filtered between 1 Hz and 7.5 kHz by the amplifier) and stored for offline analysis. A maximum of two recordings, one in layer 2/3 and another in layer 5/6, were performed in the same animal and same day before they were sacrificed. Probes were cleaned with 1% Tergazyme for at least 24 hours following the recording, then rinsed with DI water.

### In vivo experimental paradigm

In the CLA-ACC experiments, CLA somas were stimulated 150 times with 5 ms duration light pulses delivered in a 5-second inter-trial interval through an optic fiber (200 µm diameter, 5 mW at the tip of the fiber, 470 nm, Prizmatix, Israel) located above the CLA. Simultaneously, the ACC activity was recorded as described above. At the end of the protocol, the recording was stopped. If no changes in the LFP activity were detected online, the probe was moved 100-200 um deeper and then repeated the stimulation protocol. To corroborate that CLA was activated, in some experiments, 5 ms and 500 ms duration light pulses were delivered every 5 seconds at least 50 times directly into the CLA using a silicon probe attached to an optic fiber (A1xOA32LP-Poly2-5mm-50 s-177, NeuroNexus, MI, USA). If no optotagged units were observed online, the probe was lowered at a low speed (1 μm/sec) for a maximum of 200 µm and repeated the photostimulation. Our criteria for determining whether CLA was activated was a positive change in firing rate in the units in the 1-10 ms window after 5 ms photostimulation. In the 500 ms protocol, some units sustained the activity while photostimulation was evoked. Others adapted a few milliseconds after the initiation, silicon probe within the CLA when brain slices were obtained. At the end of the experiment, the virus expression was corroborated. If no CLA fibers were observed in the ACC, or if the injection was predominantly out of the CLA, the animal was removed from the study.

### Spike sorting

Recordings acquired in OpenEphys were sorted using Kilosort2.5^92^. Manual curation was performed using Phy2^93, 94^. The result from Kilosort was firstly curated by controlling and separating units into either MUA, SUA or Noise. Following the curation, the SUA was analyzed in terms of the correlogram for potential merging or splitting of units. The minimum amplitude for spikes being 125 µV. Optogenetic activation of VGLUT2 and CaMKIIa were aligned with electrophysiological recordings using event times recorded using OpenEphys and custom code. Probe trajectory was estimated from histology and Allen Brain Atlas^66^. MUA and SUA which occurred within layer 2/3 or layer 5 were selected for further analysis. The waveforms for each SUA were combined individually and an average waveform was calculated. The spike width was calculated as the duration between the maximum amplitude and the repolarization to baseline. A histogram was plotted which showed a bimodal distribution of spike widths, the spike width of 0.6 ms was chosen as the threshold and used for the separation of FS and RS. The percentage of FS and RS found in each group was calculated. The firing frequency was calculated for each SUA with 10 ms bin using a gaussian kernel (sigma=10 ms) and subsequently the z-score for each SUA was calculated using custom code for each block of trials (CHANGE-TO-MUTUAL GITHUB-WITH-ROBERTO available at: https://github.com/jofrony/SilberbergLab). The responses for each SUA were separated according to either the excitation or inhibition (50 ms following activation), or no modulation through the stimulation. The z-score was ordered according to the response within the first 50 ms following the stimulation. The modulation index was calculated as the maximum divergence within the first 50 ms. The delay following the stimulation was calculated for each group using the maximum peak of the excitation (using z-score).

### Brain slice preparation

Mice were incubated for at least 21 days post-surgery following which the mice were sacrificed and used for *ex vivo* recordings. Mice were anesthetized with isoflurane and their brains removed in ice-cold cutting solution containing the following (in mM): 2.5 KCl, 1.25 NaH_2_PO_4_, 0.5 CaCl_2_, 7.5 MgCl_2_, 10 glucose, 25 NaHCO_3_, 205 sucrose. Coronal slices, 250 μm thick, were cut (Leica VT 1000S, Wetzlar, Germany), then transferred to a 35°C water bath for 1 h in artificial CSF containing the following (in mM): 125 NaCl, 25 glucose, 25 NaHCO_3_, 2.5 KCl, 2 CaCl_2_, 1.25 NaH_2_PO_4_, 1 MgCl_2_. Slices were subsequently removed from the water bath and kept at room temperature (22 – 24°C). Slices were kept for no longer than 12 h after the brain was sliced. Slices were subsequently transferred for experiments into a bath artificial CSF solution with 10 μM of gabazine and were kept for no longer than 10 hours after the brain was sliced.

### Whole-cell patch clamp recordings

Whole-cell patch clamp recordings were obtained at 35 ± 0.5°C. Glass electrodes were pulled with a Flaming/Brown micropipette puller P-97 (Sutter Instruments) and had a resistance of 7– 9MΩ. They contained (in mM) 130 K-gluconate, 5 KCl, 10 HEPES, 4 Mg-ATP, 0.3 Na-GTP, 10 Na2-phosphocreatine and 0.3% Neurobiotin. Neurons were visualized with infrared differential interference contrast (IR-DIC) microscopy (Zeiss FS Axioskop) and fluorescent microscopy, using a mercury lamp (HBO 100, Zeiss) and a fluorescent filter cube mounted on the same microscope. Pairs of neurons were recorded within a range of 200μm from each somata. Recordings were amplified using MultiClamp 700B (Molecular Devices) and digitized by an ITC-18 (HEKA Elektronik) acquisition board. Data were acquired and analyzed using IGOR Pro (WaveMetrics).

### Determination of cortical layer

Cortical layers were assigned to every neuron prior to recordings. Layer 1 is defined as the layer closest to the pia with few neurons. Layer 2/3 is defined as a dense layer of neurons next to layer To avoid attribution errors, both infragranular layers 5 and 6 are considered together. Layer 5/6 is defined as neurons located above the cingulum bundle (Fig. 3a, b).

### Photostimulation in brain slices

Photostimulation was generated through a 1-watt blue LED (wavelength 465 nm) mounted on the microscope oculars and delivered through the objective lens. Photostimulation was controlled by a LED driver (Mightex Systems) connected to the ITC-18 acquisition board, enabling control over the duration and intensity. The photostimulation diameter through the objective lens was ∼400 μm with an illumination intensity of 16 mW/mm^2^ for experiments on VGLUT2 mice and 0.8 mW/mm^2^ for all other experiments. Short light pulses (2 ms duration, 8 pulses at 10 Hz, followed by a single 2 ms light flash after another 500 ms) were delivered to evoke postsynaptic responses in neurons; these responses were recorded with the patch pipette. Every experiment was repeated 20 times with 10 second intervals. A recording was determined to receive synaptic input if it showed repeatable stimulus-locked post-synaptic response to photostimulation.

### Confocal imaging

Some slices used for *ex vivo* whole-cell patch clamp recordings were fixed in Lana’s fixative (4% PFC with picric acid) in preparation for imaging. Slices were fixed overnight and subsequently washed in PBS 6 times, for 10 minutes each wash. A subset of slices with cells intracellularly filled with neurobiotin were next incubated in Cy5-PBS for at least 72 hours. Slices were then mounted on a glass slide with 70% glycerol and DABCO mounting medium. Confocal images were acquired using confocal microscopy (Zeiss LSM 510).

