## Supplementary Figures for "Presynaptic and Postsynaptic Determinants of the Functional Connectivity Between the Claustrum and Anterior Cingulate Cortex"

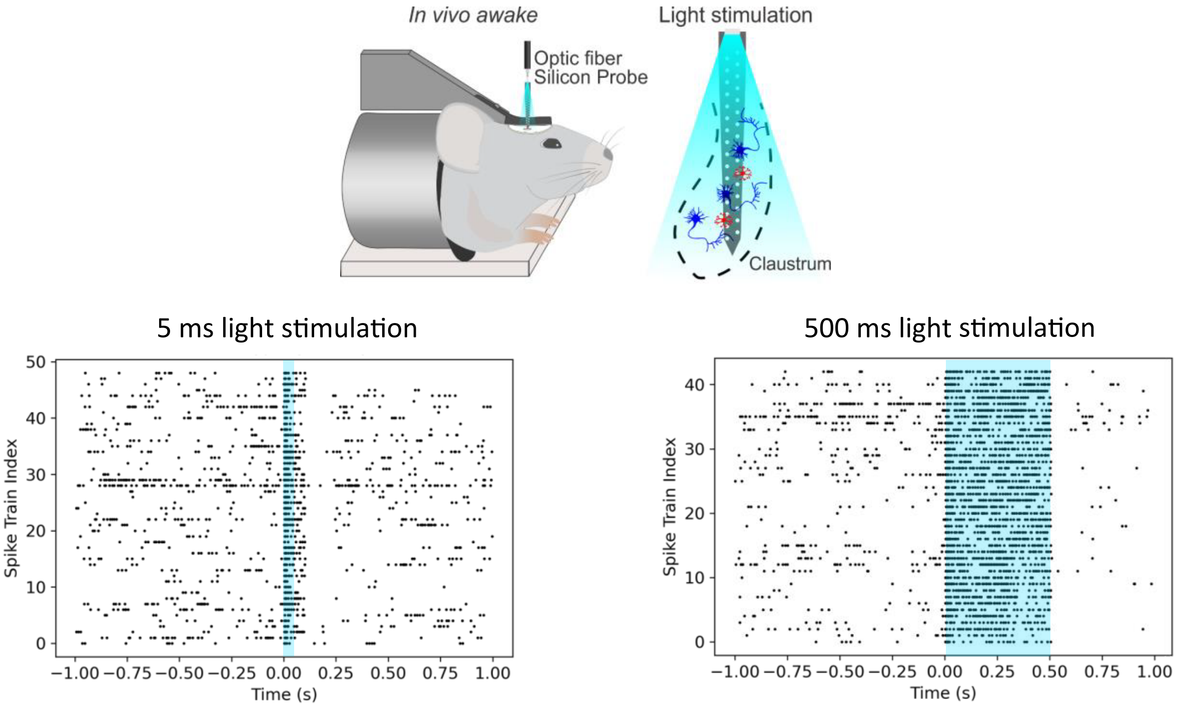


**Supplementary Fig. 1 (related to Fig. 1 and 2) Optogenetic identifiication and stimulation of VGLUT2 CLA neurons in awake head-restrained mice.**

Schematic of the in vivo recordings and photostimulation of the VGLUT2 neurons in the CLA. Raster plot showing the robust response (50 repetitions) of VGLUT2-CLA units in response to 5 ms and 500 ms light stimulation.

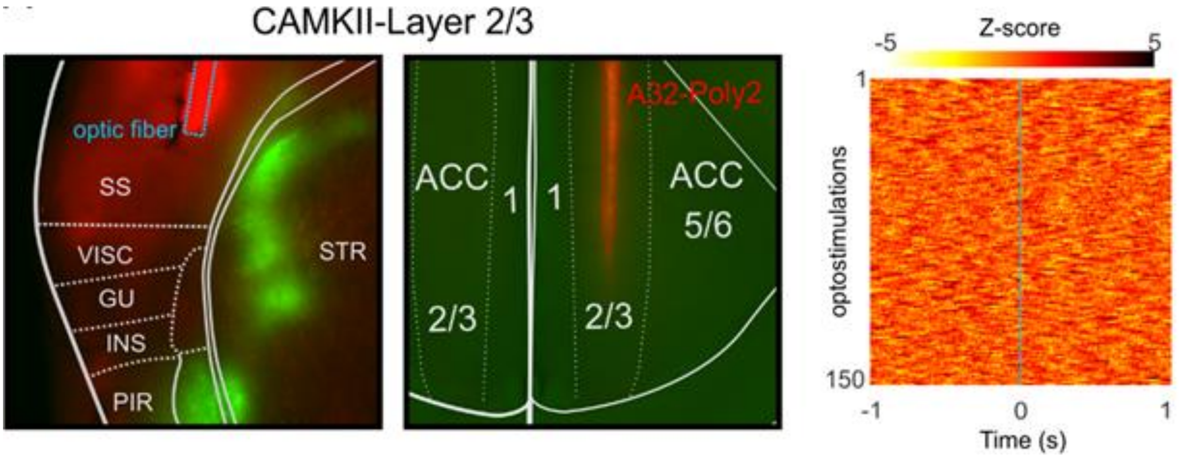


**Supplementary Fig. 2 (related to Fig. 1) ChR2+ expression outside of the CLA did not result in any functional responses or photoelectrical artifacts in the ACC following CLA photostimulation in awake head-restrained mice.**

Confocal image of a 250 µm experimental slice in a PV-tdTomato mouse, with a close-up in the CLA (left panel) and ACC region (right panel) showing virus injection outside the CLA and lack of YFP expression in the fibers in the ipsilateral ACC. In red, the optic fiber (left panel) and the silicon probe (right panel) are indicated. Heat map showing 150 repetitions with no LFP responses in the ACC layer 2/3 to 5 ms photostimulation of CLA neurons driven by a CamKIIa promoter.

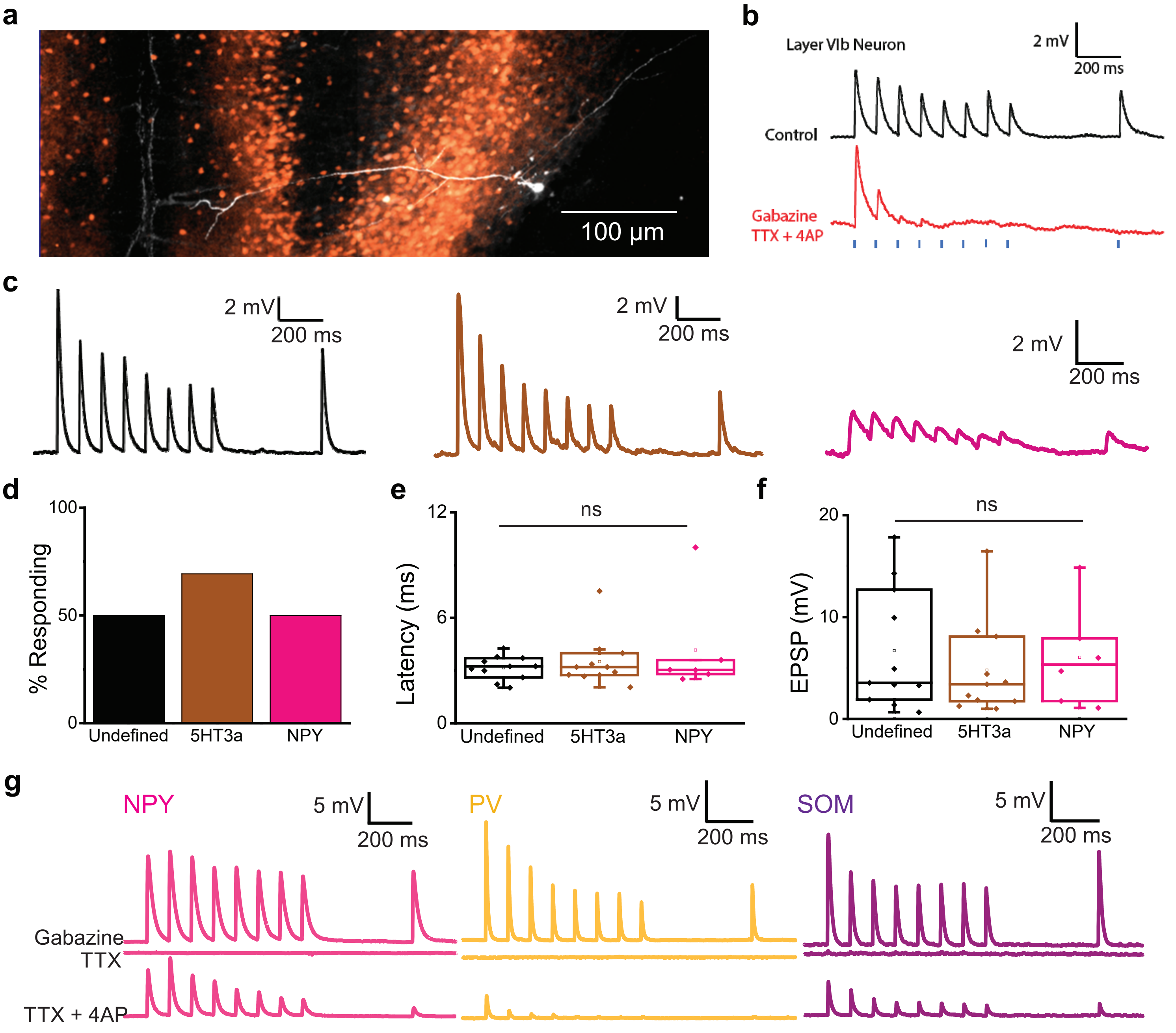


**Supplementary Fig. 3 (related to Fig. 7 and 8) Layers 1 and 6b neurons in the ACC also receive claustrum input.**

**a**. Example of neurobiotin-filled layer 6b neuron. **b**. Layer 6b neurons in the ACC received monosynaptic input from the claustrum (n = 3). **c**. Representative EPSP traces of undefined, SOM and 5HT3a layer 1 interneurons to stimulation of claustrum projections at 10Hz photostimulation rate. **d**. Percentage of Layer 1 recordings that respond to claustrum input: undefined PSNs (50%, 8/16 recordings) SOM (100% 1/1 recording), 5HT3a (66%, 8/12 recordings), NPY (20%, 1/5 recordings). **e**. No significant difference in response latency between cells in layer 1 ($P > 0.05$, One Way ANOVA). **f**. No significant difference in EPSP amplitude between cells in layer 1 ($P > 0.05$, One Way ANOVA). **g**. Postsynaptic responses from different interneuron types are recorded in gabazine. The postsynaptic response is subsequently removed on bath application of TTX and recovered with TTX+4AP, this confirms that the input is monosynaptic.
